# CRitical Assessment of genomic COntamination detection at several Taxonomic ranks (CRACOT)

**DOI:** 10.1101/2022.11.14.516442

**Authors:** Luc Cornet, Valérian Lupo, Stéphane Declerck, Denis Baurain

## Abstract

**Background:** Genome contamination is a well-known issue in (meta)genomics. Although it has received a lot of attention, with an increasing number of detection tools made available over the years, no comparison between these tools exists in the literature.

**Results:** Here, we report the benchmarking of six of the most popular tools using a simulated framework. Our simulations were conducted on six different taxonomic ranks, from phylum to species. The analysis of the estimated contamination levels indicates that the precision of the tools is not good, often due to large overdetection but also underdetection, especially at the genus and species ranks. Furthermore, our results show that only redundant contamination is accurately estimated.

**Conclusion:** Our results indicate that using a combination of tools, including Kraken2, is necessary to estimate the contamination level accurately. We also provide a freely available contamination simulation framework, CRACOT, which may be useful for estimating the accuracy of future algorithms.

## Background

Genomic contamination is a well-known, albeit recurrent, problem in genomics. It appears when a genome, often a Metagenome-Assembled Genome (MAG), contains DNA sequences that do not belong to the expected organism [1]. This umbrella concept actually masks different sources of DNA mis-affiliation, which can occur almost anytime between the selection of a sample and its bioinformatic analysis [1]. Nowadays, genomes are the basis of numerous studies, and it is no longer necessary to demonstrate that genomic contamination is a cause for artifacts, notably in phylogenomic inference [2] [3] [4]. Consequently, the detection of contaminants is a topic that has attracted the attention of scientists, with the development of numerous detection tools and an increasing rate of publications over the recent years. Although all these tools ultimately report a quantified level of contamination, they are based on various algorithms and do not measure the same information [1] [5]. Indeed, among the most popular tools, two major categories can be distinguished: those relying on the presence of multiple marker genes (e.g., CheckM [6] and BUSCO [7]) and those based on whole-genome surveys (e.g., GUNC [8], Physeter [5] and Kraken2 [9]). Because of these differences in algorithms, Cornet et al 2018 [10] and Lupo et al 2021 [5] have reported the difficulty to meaningfully compare these tools, let alone computing correlations between their estimates. Simulations of genome contamination, in which the exact amount of contaminant sequences is known, can nevertheless be used to overcome such a limitation. In the present study, we compare the detection performance of six of the most used tools (CheckM [6], BUSCO [7], GUNC [8], Physeter [5], Kraken2 [9], and CheckM2 [11]) in order to assess their efficiency. To do so, we used simulations at multiple taxonomic ranks, while varying the contamination scenarios.

## Methods

### Contamination simulations (overview of CRACOT)

Our genome contamination simulations were carried out with the Nextflow workflow CRACOT, freely available at https://github.com/Lcornet/GENERA/wiki/20.-CRACOT.

705 high quality genomes belonging to class *Clostridia* and genus *Lactobacillus* were selected as input for CRACOT. These genomes were selected based on the GUNC [8] clade separation score (CSS), which measures the chimerism of genome contigs. Furthermore, we imposed on these genomes to have no more than five contigs with no ‘N’ within each contig. The contamination values of these genomes for the six tools are available in **Table S1**. The median contamination was 0.45% for CheckM V1.2.1 [6], 0.02% for GUNC V1.0.5 [8], 0.87% for BUSCO V5.4.3 [7], 22,3% for Physeter V0.213470 [5], 2,45% for Kraken2 V2.1.2 [9], 8% for CheckM2 V0.1.3 [11].

The first step of CRACOT, **Figure 1**, was to create random genome pairs, one genome being considered hereafter as the main “expected” organism and the second as the slave “contaminant” organism. The pairing, based on the NCBI Taxonomy [12] [13] provided by Bio-Must-Core V0212670 (https://metacpan.org/dist/Bio-MUST-Core), associated with the genomes was made for one specific taxonomic rank, ranging from phylum to species. When a rank is selected, the two genomes should belong to the same taxon at this rank, but have a different taxonomy starting with the next rank. For instance, the phylum rank (e.g., Firmicutes) means that the two genomes belong to the same phylum but not to the same class (e.g., if one genome belongs to Bacilli, the other genome belongs to Clostridia).

**Figure 1:**
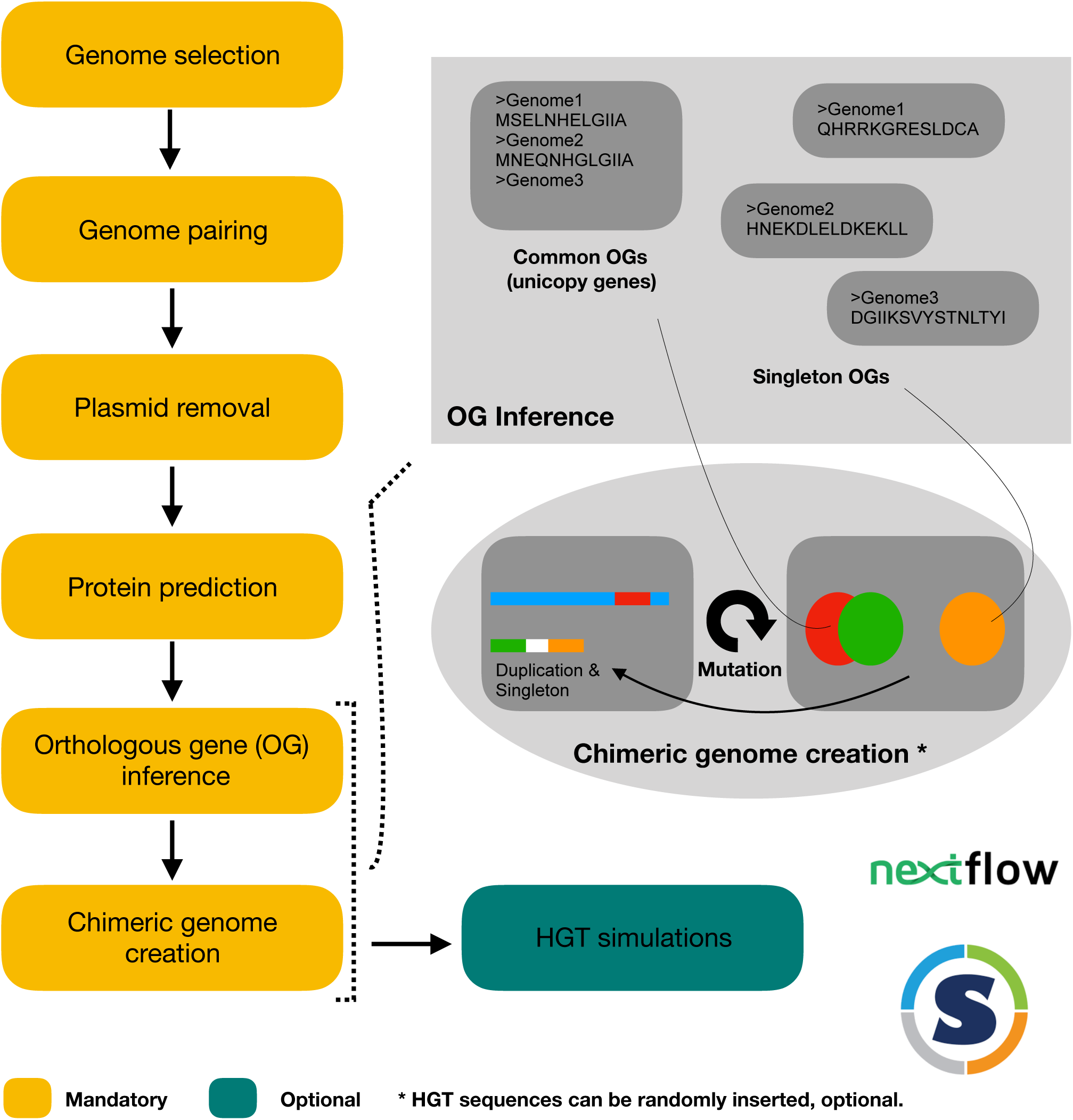
Flowchart of CRACOT. CRACOT is a Nextflow workflow, supported by a Singularity container. It is a six step program. The first step is the genome selection according to a user-specified list. The second step is the association of genomes, by pairs of the same taxonomic group. Step 3 to 5 corresponds to the removal of plasmid, protein prediction and orthology inference. Finally, genome contamination simulations are based on the information produced during the orthology inference, the common (to both the expected and contaminant organisms) and single (of only the contaminant organism). Common genes are used for redundant and replaced contamination events while singleton are used for single contamination events. Optionally, a mutation rate can be enabled for each of these three basic types to simulate horizontal gene transfer.

The plasmids of the selected genomes were removed after the pairing step, so as to not interfere with the detection of contamination. Removal was performed with PlasmidPicker (https://github.com/haradama/PlasmidPicker) with default settings. Proteins werethen predicted with Prodigal V2.6.3 [14], used with default settings. Finally, OrthoFinder V2.5.4 [15], with default settings, wasused for orthologous inference.

The three types of contamination were simulated based on the common and single protein orthogroups (OGs). Common proteins weredefined as proteins present in only one copy for both the main and slave genome in the OG while single proteins were singletons of the slave genome. Duplicated contamination events were fished from the pool of common OGs, and the corresponding gene sequences of the slave genome were added to the end of the last contig of the master genome (with a serie of five ‘N’ added to either side of the gene). Replaced contamination events were also fished from the pool of common OGs but slave genes replaced the genuine genes within the main genome. Single contamination events were fished from the pool of singletons of the slave organism, and the corresponding gene sequences were added to the end of the last contig of the master genome, as above. The number of events of each type is a user-specified option. At each simulation, 150 chimeric genomes were asked as CRACOT output, but the real output number depends on the number of available common and single protein OGs. The number of simulated genomes used in this study are given in **Table S2**. Chimeric levels of the simulations are indicated in **Table S3**.

HGT can be simulated for each of the three sequence types. The mutation rate was computed with HgtSIM [16], with the rate option set at 1-0-1-1 so that a mutation rate in DNA sequences corresponds to the same simulation rate in the proteins [16].

CRACOT was used to simulate contamination events, not only the redundant, replaced or single type separately, but also as a combination of the three types. Two HGT simulations for a combination of the three contamination types, with a mutation rate of 10% and 25%, were also generated.

### Genomic contamination estimation

Genomic contaminants were estimated using the Nextflow workflow GENcontams (https://github.com/Lcornet/GENERA/wiki/09.-Genome-quality-assessment) from the GENERA project [17]. CheckM V1.2.1 [6] was used with the lineage_wf option and the provided database. GUNC V1.0.5 [8] was used with default settings and the Progenomes 2.1 database [18]. BUSCO V5.4.3 [7] was used in auto-lineage mode and the provided database. BUSCO’s number of duplicated markers was used as a proxy for the contamination level. Physeter V0.213470 was used with the auto-detect option and the database provided in Lupo et al. (2021) [5]. Kraken 2 V2.1.2 [9] was used with default settings and the database ‘PlusFP’ downloaded from https://benlangmead.github.io/aws-indexes/k2. Kraken2 levels of contamination were computed with the Physeter parser with the auto-detect option set to ‘count_first’. The list of taxa used by the Physeter parser was automatically produced by the create-labeler.pl script using the list of genera found in the nodes.dmp file from the local mirror of NCBI Taxonomy. CheckM2 V0.1.3 [11] was used with default settings and the provided database.

### Correlation and violin plot creation

Spearman correlations between the CL estimates of the tools and the simulated levels of contaminants, as created by CRACOT, were computed with R [19]. Violin plots were created with ggplot [20]. The R code for the creation of these plots is available at https://github.com/Lcornet/GENERA/wiki/21.-Supplemental-Scripts#4-make-cracot-tablepy-and-cracot-rscript.

## Results and Discussion

Regardless of the contaminant source, it is now established that it can be summarized into three main types at the genomic sequence level (**Figure 1**) [1] [8]. The first type is **redundant contamination** that occurs when the contaminant sequence is redundant with an homologous genomic sequence of the expected organism [1]. The second type is **replaced contamination** that is similar to the first one, but with the genuine sequence of the expected organism lacking from its genome [1]. The third type is **single contamination** that occurs when the contaminant sequence has naturally no homologous sequence within the genome of the expected organism [1]. To mimic these three situations, we selected 705 high-quality reference genomes belonging to class *Clostridia* and genus *Lactobacillus* and simulated contamination events of the three types (**Figure 1**). Our simulations were performed out at six different taxonomic ranks, from intra-phylum to intra-species.

Surprisingly, our results reveal that, with the exception of Kraken2, none of the tested tools was able to accurately estimate the contamination level (CL) of our combined scenarios, when the three different contamination types were mixed (**Figure 2**). Separated simulations are available in Supplementary materials for redundant (**Figure S1**), replaced (**Figure S2**), and single (**Figure S3**) events. CheckM, based on the duplication of gene markers [6], overestimated the redundant CL (**Figure S1**), but quite logically, does not detect replaced (**Figure S2**) or single (**Figure S3**) contamination events. Similar to its main metric, CheckM’s complementary metric used for genetically close contaminants (strain heterogeneity) also overestimated CL, but at the genus and species ranks (**Figure 2**). BUSCO, which is also based on marker duplication [7], largely overestimated the redundant CL (**Figure S1**) at all ranks and, as for CheckM, underdetected replaced (**Figure S2**) and single (**Figure S3**) contamination events. GUNC, which searches for sequence chimerism [8], presents a pattern of both over- and underestimation at four ranks (phylum, class, order and family) (**Figure 2**), with a minimum of 59% of underestimation (see **Table S4** for the percentage of underestimation of each tool at each taxonomic rank). At the genus and species ranks, GUNC only underestimated CL (**Figure 2**), notably for replaced events where it detected nothing (**Figure S2**). Physeter, which is based on Lowest Common Inference (LCA) of DIAMOND blastx [21] hits [5], overestimates CL at all ranks for all types of contaminants (**Figure 2**). In contrast, Kraken2, which takes advantage of exact long kmer matching [9], showed the best estimation of CL, fitting well to the simulations, with the exception of the species rank, which was largely underestimated (see **Table S4**). It is noteworthy that the genomes used in our simulations were of high quality (see Online Methods) and included in the Kraken2 database. Owing to its exact kmer matching algorithm [22], one cannot exclude that Kraken2 would perform less well on rare genomes, compared to our simulations. CheckM2, which uses machine learning based on genomic contamination simulation (gradient boost model) without relying on taxonomic information, [11], largely overestimated the redundant CL (**Figure S1**), especially at the genus and species ranks. Replaced (**Figure S2**) and single (**Figure S3**) CL were underestimated at all ranks, with the exception of the single type at the genus and species ranks. Percentages of underestimation(**Table S4)** showed that CheckM2 underestimates CL in more than 97% of the cases for both replacement and single events, while it never underdetected the redundant type (to the exception of the species rank in 1.3% of the cases). To overcome the impossibility to directly correlate the performance of the different tools (due to their algorithmic differences), we computed the correlation of each tool to the expected CL of our simulations. The tools, not including Kraken2, correlated badly, often negatively, with the expected CL level of the simulations, the correlation coefficient (R2) never going beyond 0.37 (**Figure 2, Figures S1-S3**).

**Figure 2:**
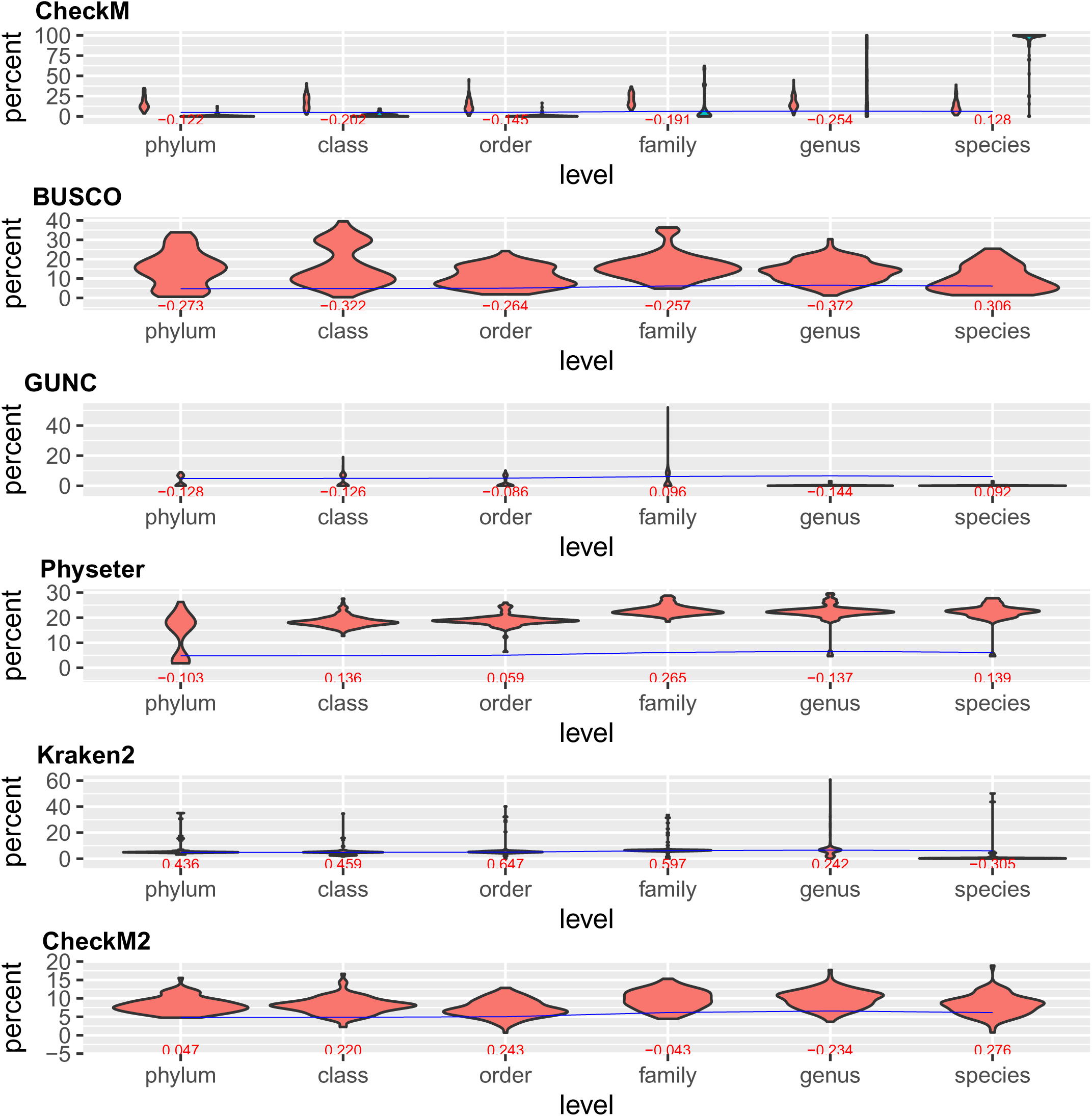
Contamination estimation, at six taxonomic ranks, of the combined types of contamination. Simulations were performed with a combination of the three contamination types (redundant, replaced, single). The median values of the contamination level (%CL) of these simulations are indicated by the blue line, while these CL estimated by the six tools are summarized by the violin plots. Spearman correlation values between the estimate of each tool and the simulated level of contamination are indicated in red.

Beside genomic contamination, another kind of genomic exchange naturally affects genomes: horizontal gene transfer (HGT). One of the major differences between HGT and contamination is that the first one accumulates mutations in the receiver (and donor) organisms [23] after transfer whereas contamination occurs shortly before or after genome sequencing, hence contaminant sequences are exact matches between donor and receiver genomes [1]. To investigate the effect of HGT on detection performance, a non-null mutation rate was optionally enabled during the simulations (see Online Methods), either at 10% (**Figure S4**) or 25% (**Figure S5**). None of the tools (to the exception of Physeter) conflated contamination and HGT, which suggested that HGT events should not increase CL on real data. While reassuring, a possible drawback is that if the contaminant sequence is an HGT, it has few chances to be detected. This can be damaging since HGT frequently occurs in bacteria [24] [25] [26] [27] [28]. Somewhat ironically, the inability of Physeter to differentiate between HGT and genomic contamination indicates that LCA algorithms are likely to prove useful in this case, even if too conservative due to their inclination for overdetection.

## Conclusion

We conducted this study because no tools comparison, despite the availability of no less than 18 programs, had been published to date, raising the question in the community: “which tool should we use?” Our results have demonstrated that CL is frequently overestimated, resulting in unwarranted removal of sometimes precious genomes. Nevertheless, especially at the genus and species ranks, the odds of underestimation are always significant. This is a matter of concern because the risk of contamination by closely related taxa is higher when dealing with MAGs. We have also demonstrated that the replaced and single contamination types suffer less from underestimation compared to the redundant events. The results of this study are all the more surprising asour simulations were rather simple. Furthermore, simulations were conducted with only one contaminant genome, at low CL, while contamination by more than one taxon, at high CL, regularly occurs in public repositories [10] [5]. Our conclusion is that, given the current algorithmic state of the field, which requires more innovation, users should use a combination of tools to estimate CL, and one of these tools should be Kraken2. Our contamination simulation framework, CRACOT, is freely available as a Nextflow workflow [29], sustained by a Singularity container [30], at https://github.com/Lcornet/GENERA/wiki/20.-CRACOT. It might be useful in future projects, for example to estimate the accuracy of new tools underdevelopment.

## Abbreviations

(MAG): Metagenome-Assembled Genome
(CL): contamination level
(HGT): horizontal gene transfer
(CSS): clade separation score
(OGs): orthogroups

## Declarations

### Ethics approval and consent to participate

Not applicable.

### Consent for publication

Not applicable.

### Availability of data and materials

CRACOT is freely available at https://github.com/Lcornet/GENERA/wiki/20.-CRACOT.

### Authors’ contributions

LC and DB conceived the study. LC developed CRACOT and performed all analyses. VL developed the parser of Kraken2, used for CL estimation. LC and DB wrote the manuscript with the help of VL and SD.

## Acknowledgements

Not applicable.

