## Supplemental data for "CRitical Assessment of genomic COntamination detection at several Taxonomic ranks (CRACOT)"

### Supplemental file 1

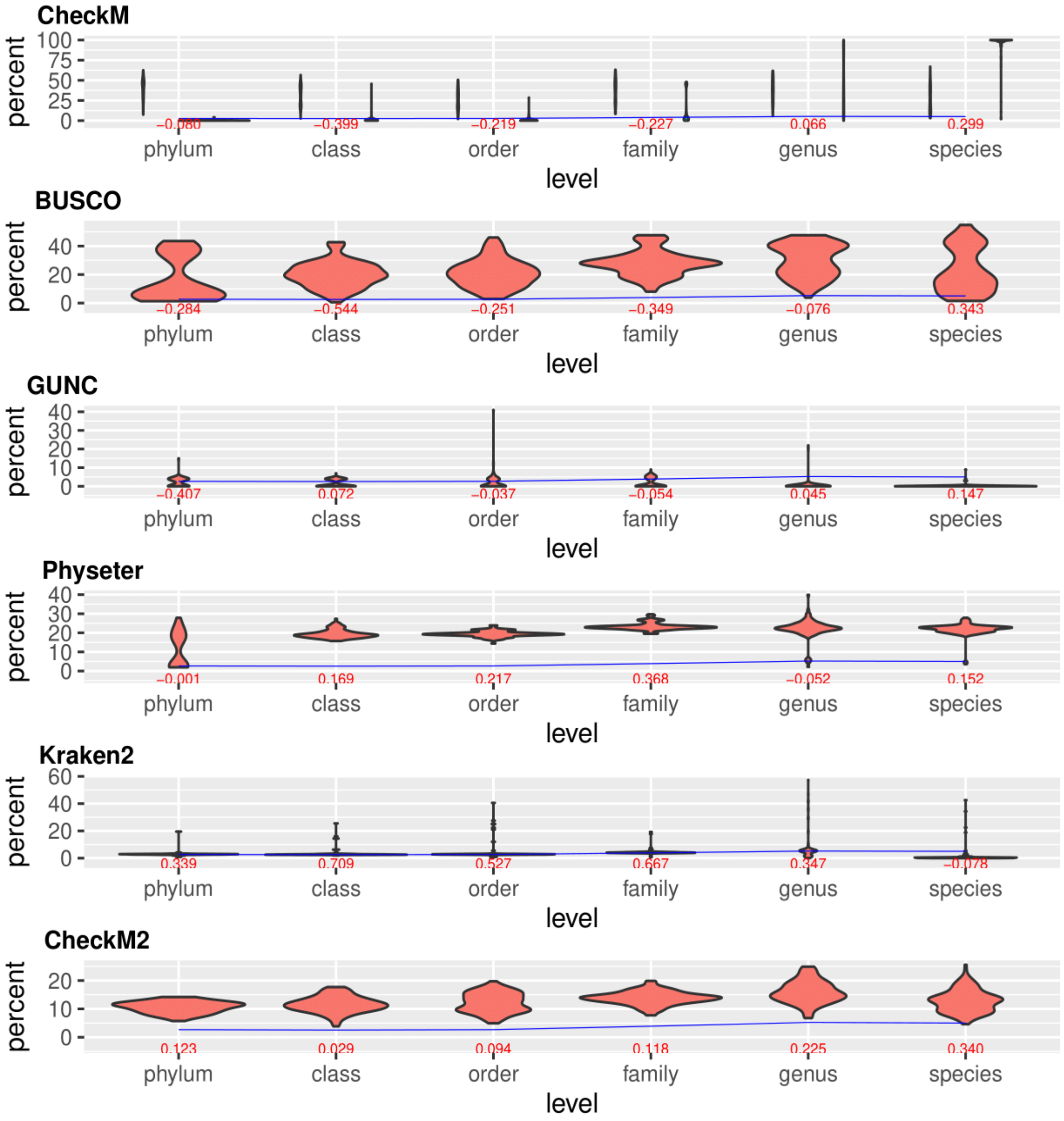


**Figure S1: Contamination estimation, at six taxonomic ranks, of the redundant type of contamination.**

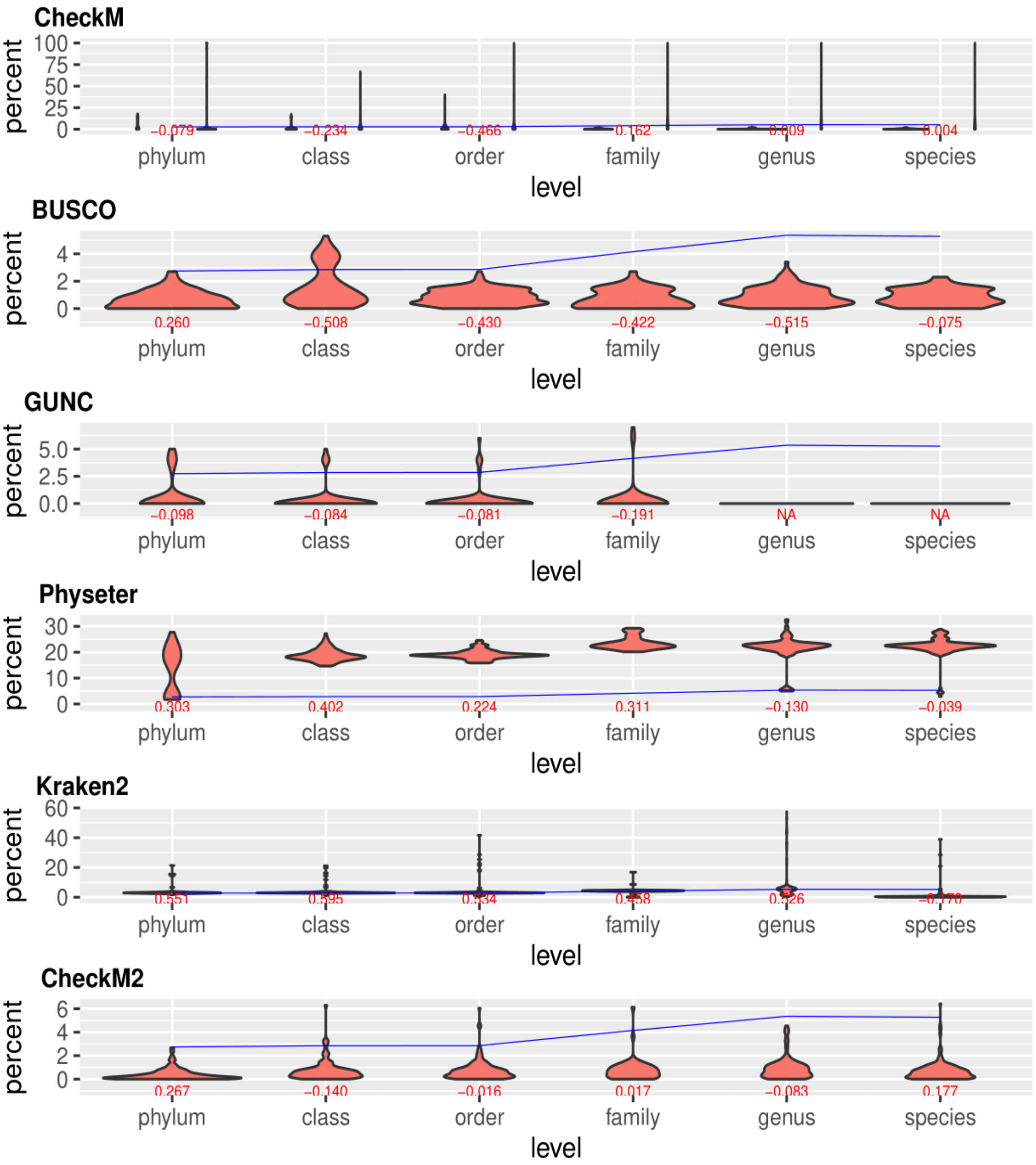


**Figure S2: Contamination estimation, at six taxonomic ranks, of the replaced type of contamination.**

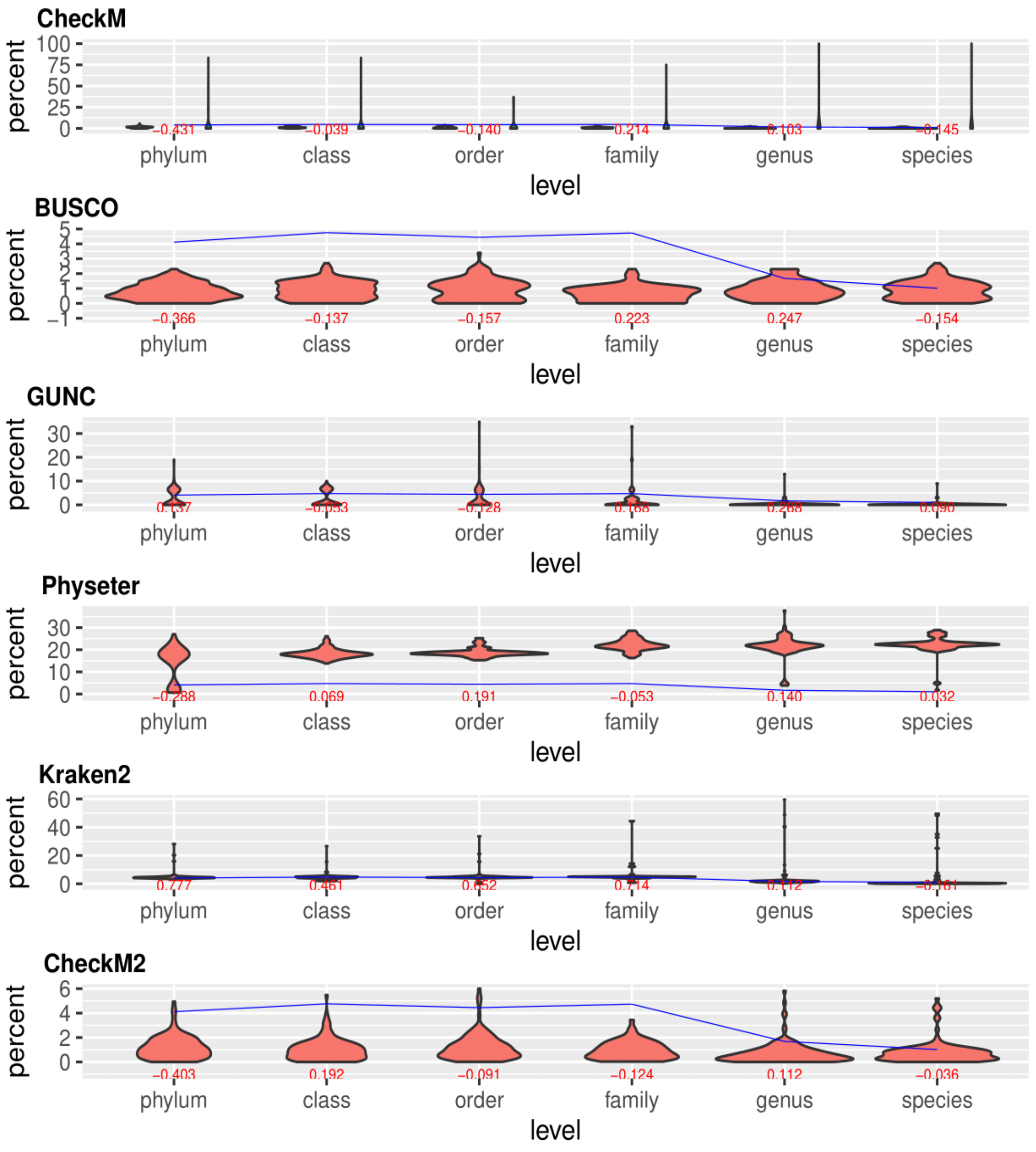


**Figure S3: Contamination estimation, at six taxonomic ranks, of the single type of contamination.**

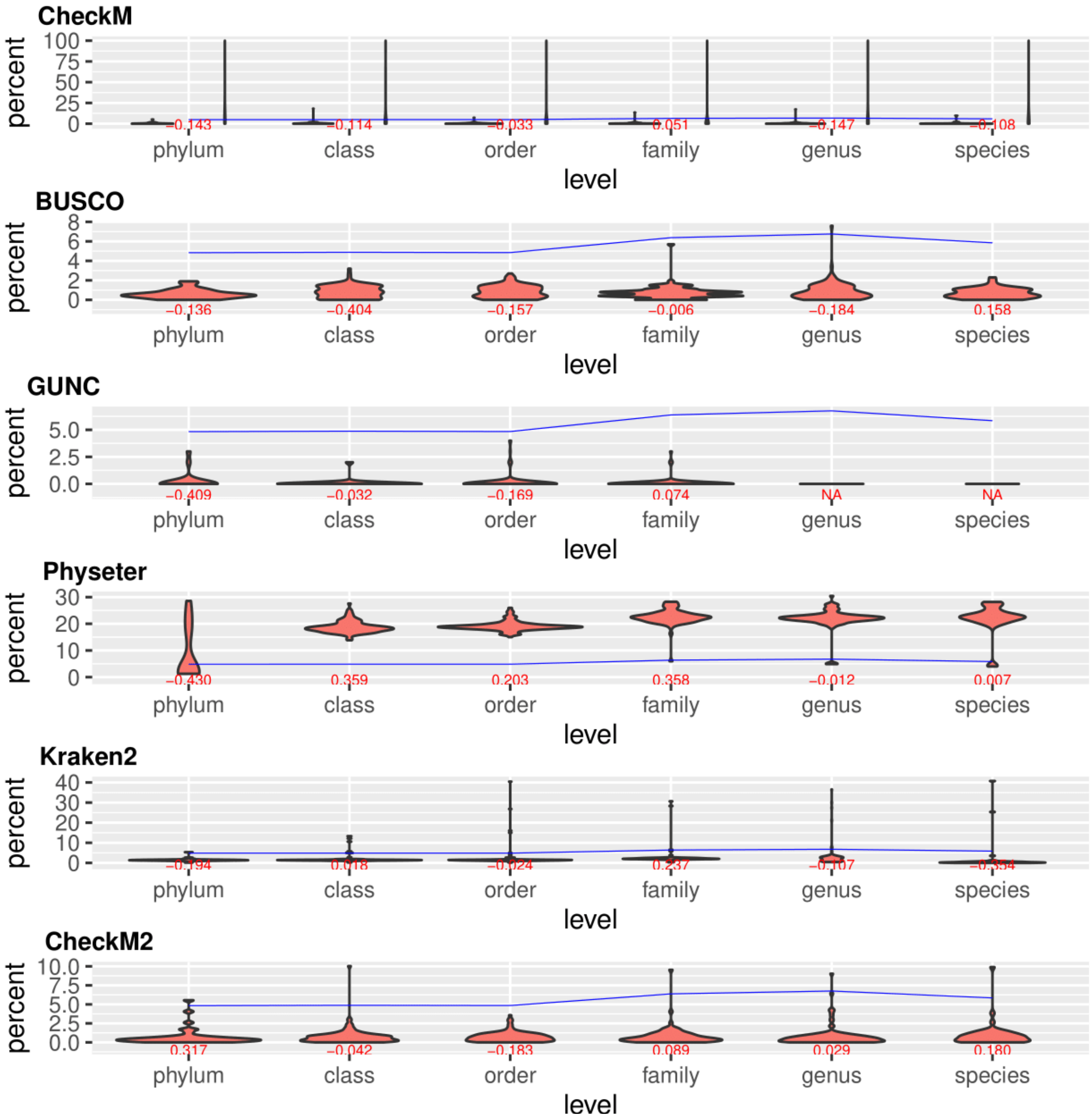


**Figure S4: Contamination estimation, at six taxonomic ranks and with a mutation rate of 10%, of the combined types of contamination.**

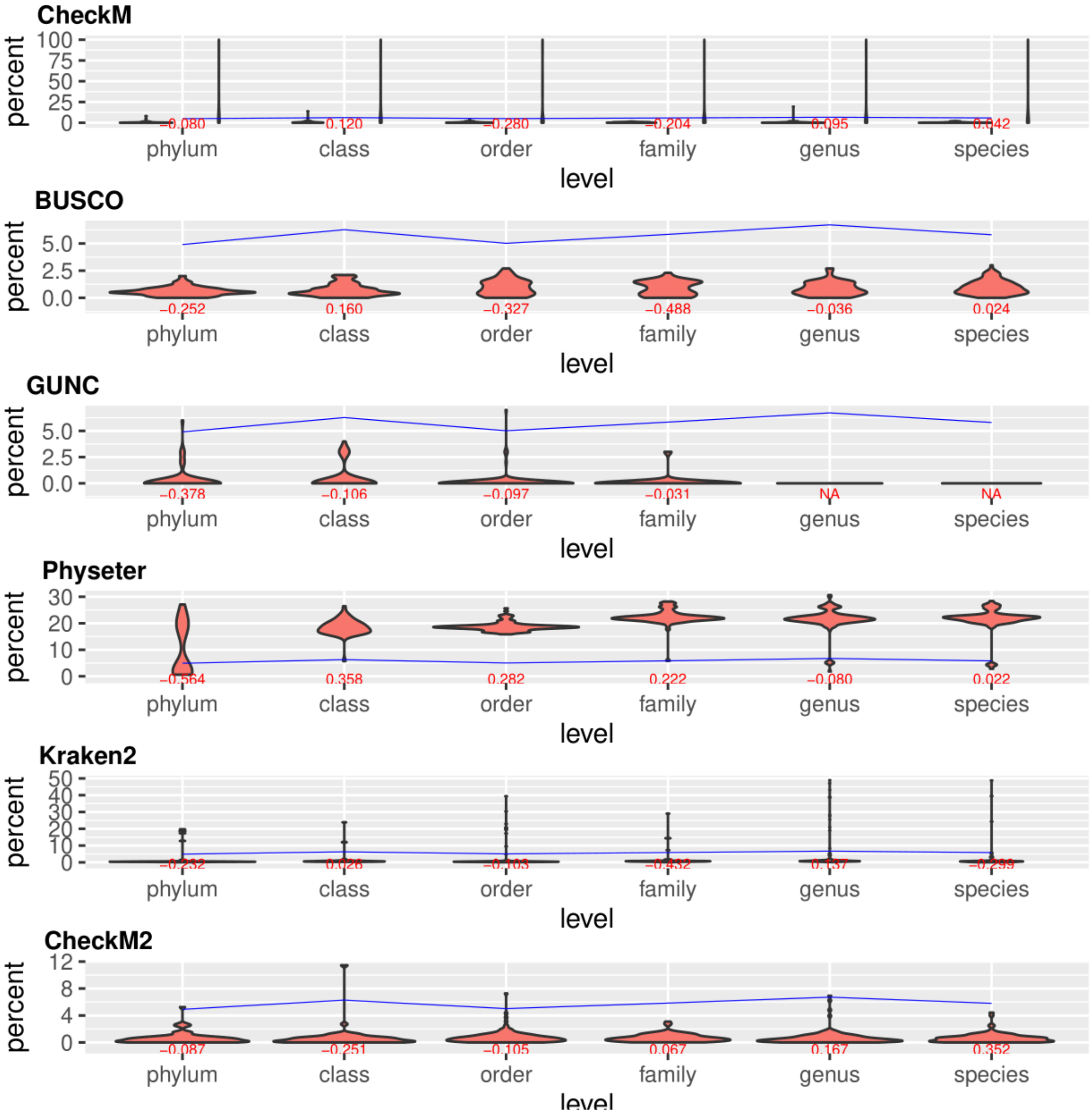


**Figure S5: Contamination estimation, at six taxonomic ranks and with a mutation rate of 25%, of the combined types of contamination.**

| Genome | Checkm | Busco | GUNC | Physeter | Kraken | Checkm2 |
|---|---|---|---|---|---|---|
| GCF_902388275.1 | 0.00 | 1.1 | 0.0 | 24.76 | 0.14 | 10.09 |
| GCF_902386435.1 | 0.00 | 0.4 | 0.0 | 22.63 | 0.51 | 10.13 |
| GCF_901523095.1 | 0.00 | 0.4 | 0.0 | 22.98 | 0.35 | 0.06 |
| GCF_901482515.1 | 0.00 | 0.4 | 0.0 | 22.95 | 0.25 | 0.04 |
| GCF_900478015.1 | 0.00 | 0.8 | 0.0 | 22.73 | 0.44 | 0.07 |
| GCF_900461415.1 | 0.00 | 0.4 | 0.0 | 23.21 | 0.34 | 0.53 |
| GCF_900461395.1 | 0.00 | 0.4 | 0.0 | 22.88 | 0.98 | 0.29 |
| GCF_900461135.1 | 0.00 | 0.4 | 0.0 | 22.59 | 0.51 | 0.34 |
| GCF_900447135.1 | 0.00 | 0.8 | 0.0 | 22.12 | 0.54 | 10.13 |
| GCF_900447095.1 | 0.00 | 0.4 | 0.0 | 23.10 | 0.34 | 10.13 |
| GCF_900447085.1 | 0.00 | 0.8 | 0.0 | 23.15 | 1.15 | 0.1 |
| GCF_900447035.1 | 0.00 | 0.4 | 0.0 | 23.49 | 0.41 | 0.43 |
| GCF_900209925.1 | 1.34 | 0.0 | 0.0 | 22.27 | 0.18 | 10.61 |
| GCF_900184165.1 | 0.24 | 0.0 | 0.0 | 21.42 | 0.98 | 0.04 |
| GCF_021561035.1 | 0.70 | 1.5 | 0.0 | 22.37 | 12.68 | 0.29 |
| GCF_021560735.1 | 0.00 | 0.8 | 0.0 | 21.97 | 0.46 | 0.14 |
| GCF_020809405.1 | 0.00 | 1.5 | 0.0 | 22.28 | 0.13 | 6.01 |
| GCF_020139295.1 | 0.00 | 0.4 | 0.0 | 22.69 | 0.35 | 0.4 |
| GCF_020139 | 0.00 | 0.8 | 0.0 | 21.31 | 0.26 | 0.25 |

| | | | | | | |
|---|---|---|---|---|---|---|
| 255.1 | | | | | | |
| GCF_020139 235.1 | 0.00 | 0.8 | 0.0 | 22.16 | 0.44 | 0.07 |
| GCF_020139 215.1 | 0.00 | 0.8 | 0.0 | 22.45 | 0.45 | 10.13 |
| GCF_020138 795.1 | 0.00 | 0.8 | 0.0 | 22.36 | 0.40 | 0.09 |
| GCF_020138 775.1 | 0.00 | 0.8 | 0.0 | 23.14 | 0.30 | 0.14 |
| GCF_020138 755.1 | 0.00 | 0.4 | 0.0 | 23.28 | 0.31 | 0.37 |
| GCF_020138 735.1 | 0.00 | 0.4 | 0.0 | 21.64 | 0.39 | 0.68 |
| GCF_020138 715.1 | 0.00 | 0.4 | 0.0 | 22.04 | 0.38 | 0.17 |
| GCF_020138 695.1 | 0.00 | 0.4 | 0.0 | 22.36 | 0.25 | 10.13 |
| GCF_020138 675.1 | 0.00 | 0.4 | 0.0 | 23.23 | 0.30 | 0.39 |
| GCF_020138 655.1 | 0.00 | 0.8 | 0.0 | 22.31 | 0.29 | 0.67 |
| GCF_020138 635.1 | 0.00 | 0.4 | 0.0 | 22.54 | 0.27 | 0.47 |
| GCF_020138 595.1 | 0.00 | 0.4 | 0.0 | 22.24 | 0.46 | 0.07 |
| GCF_020138 575.1 | 0.00 | 0.4 | 0.0 | 22.62 | 0.23 | 0.08 |
| GCF_020138 555.1 | 0.00 | 0.4 | 0.0 | 23.04 | 0.27 | 0.09 |
| GCF_020138 535.1 | 0.00 | 0.4 | 0.0 | 22.67 | 0.38 | 0.3 |
| GCF_020138 515.1 | 0.00 | 0.4 | 0.0 | 21.93 | 0.41 | 0.45 |
| GCF_020138 475.1 | 0.00 | 0.4 | 0.0 | 21.96 | 0.41 | 10.13 |
| GCF_020138 455.1 | 0.00 | 0.4 | 0.0 | 21.95 | 0.49 | 0.06 |
| GCF_020138 | 0.00 | 0.4 | 0.0 | 21.57 | 0.36 | 0.31 |

| | | | | | | |
|---|---|---|---|---|---|---|
| 375.1 | | | | | | |
| GCF_020138355.1 | 0.00 | 0.4 | 0.0 | 22.33 | 0.46 | 0.06 |
| GCF_020138335.1 | 0.00 | 0.4 | 0.0 | 22.33 | 0.31 | 0.13 |
| GCF_019977655.1 | 0.00 | 1.9 | 0.0 | 18.81 | 0.06 | 1.31 |
| GCF_019969455.1 | 0.00 | 0.8 | 0.0 | 22.35 | 3.36 | 12.06 |
| GCF_019969425.1 | 0.00 | 0.8 | 0.0 | 23.65 | 0.05 | 0.94 |
| GCF_019969405.1 | 0.34 | 1.1 | 0.0 | 23.10 | 3.27 | 1.33 |
| GCF_019969385.1 | 0.00 | 0.0 | 0.0 | 24.53 | 1.20 | 0.11 |
| GCF_019968075.1 | 0.00 | 0.0 | 0.0 | 25.84 | 0.05 | 0.13 |
| GCF_019968055.1 | 0.00 | 0.0 | 0.0 | 26.11 | 0.04 | 0.23 |
| GCF_019968035.1 | 0.00 | 0.8 | 0.0 | 23.71 | 0.05 | 9.98 |
| GCF_019968015.1 | 0.00 | 0.8 | 0.0 | 23.87 | 0.04 | 0.93 |
| GCF_019967995.1 | 0.00 | 0.8 | 0.0 | 23.37 | 0.04 | 0.9 |
| GCF_019967975.1 | 0.00 | 0.8 | 0.0 | 23.26 | 0.08 | 1.01 |
| GCF_019967955.1 | 0.00 | 1.1 | 0.0 | 24.89 | 0.13 | 1.36 |
| GCF_018223625.1 | 0.13 | 0.0 | 0.0 | 23.56 | 0.07 | 2.92 |
| GCF_017897335.1 | 0.00 | 0.4 | 0.0 | 23.26 | 0.35 | 0.37 |
| GCF_017352215.1 | 0.70 | 0.4 | 0.0 | 28.34 | 0.00 | 0.04 |
| GCF_017347585.1 | 0.24 | 0.2 | 0.0 | 24.15 | 0.36 | 0.04 |
| GCF_017100 | 0.69 | 0.4 | 0.0 | 22.66 | 0.27 | 10.54 |

| | | | | | | |
|---|---|---|---|---|---|---|
| 085.1 | | | | | | |
| GCF_016694795.1 | 0.00 | 1.1 | 0.0 | 24.16 | 0.21 | 0.67 |
| GCF_016027155.1 | 0.00 | 0.8 | 0.0 | 22.30 | 0.50 | 0.05 |
| GCF_016026675.1 | 0.00 | 0.4 | 0.0 | 22.18 | 0.40 | 10.13 |
| GCF_013149375.1 | 0.00 | 1.1 | 0.0 | 20.76 | 0.05 | 11.19 |
| GCF_011065855.2 | 0.20 | 0.8 | 0.0 | 20.64 | 4.96 | 1.89 |
| GCF_010509575.1 | 0.00 | 0.0 | 0.0 | 23.58 | 0.19 | 10.09 |
| GCF_008151785.1 | 0.00 | 1.1 | 0.0 | 22.99 | 0.25 | 9.42 |
| GCF_006351925.1 | 0.96 | 1.5 | 0.0 | 23.44 | 0.02 | 10.91 |
| GCF_003403315.2 | 0.00 | 0.0 | 0.0 | 22.20 | 0.11 | 11.63 |
| GCF_003350945.1 | 0.00 | 0.4 | 0.0 | 23.01 | 0.37 | 10.13 |
| GCF_003312465.1 | 0.00 | 1.1 | 0.0 | 24.61 | 0.06 | 9.98 |
| GCF_003293635.1 | 0.00 | 1.1 | 0.0 | 24.76 | 0.14 | 10.09 |
| GCF_003203455.1 | 0.00 | 0.8 | 0.0 | 20.88 | 0.66 | 10.61 |
| GCF_002865995.1 | 1.63 | 1.1 | 0.0 | 22.69 | 0.17 | 10.63 |
| GCF_002844395.1 | 0.67 | 0.8 | 0.0 | 18.99 | 0.22 | 10.7 |
| GCF_002796935.1 | 1.03 | 1.1 | 0.0 | 20.57 | 39.39 | 13.86 |
| GCF_002734145.1 | 0.00 | 0.0 | 0.0 | 23.91 | 0.17 | 10.09 |
| GCF_002586945.1 | 0.00 | 1.1 | 0.0 | 25.02 | 0.03 | 9.98 |
| GCF_002564 | 0.00 | 0.0 | 0.0 | 21.73 | 1.11 | 10.56 |

| | | | | | | |
|---|---|---|---|---|---|---|
| 225.1 | | | | | | |
| GCF_002355 795.1 | 0.00 | 0.8 | 0.0 | 23.37 | 0.30 | 10.13 |
| GCF_001950 115.1 | 0.93 | 0.8 | 0.0 | 22.89 | 1.51 | 10.63 |
| GCF_001889 325.1 | 0.69 | 1.1 | 0.0 | 21.57 | 7.65 | 11.19 |
| GCF_001856 695.1 | 0.08 | 1.1 | 0.0 | 23.09 | 0.11 | 11.19 |
| GCF_001856 645.1 | 0.08 | 1.1 | 0.0 | 23.07 | 0.14 | 11.19 |
| GCF_001854 085.1 | 0.00 | 0.4 | 0.0 | 22.63 | 0.51 | 10.13 |
| GCF_001735 765.2 | 0.00 | 0.4 | 0.0 | 21.41 | 0.37 | 10.59 |
| GCF_001705 235.1 | 1.03 | 1.1 | 0.0 | 20.57 | 39.39 | 13.86 |
| GCF_000828 305.1 | 0.16 | 0.4 | 0.0 | 22.49 | 3.44 | 10.89 |
| GCF_000807 255.1 | 0.08 | 1.1 | 0.0 | 23.06 | 0.15 | 11.19 |
| GCF_000807 175.1 | 0.08 | 1.1 | 0.0 | 23.23 | 0.15 | 11.19 |
| GCF_000789 355.1 | 0.00 | 0.4 | 0.0 | 20.92 | 0.15 | 10.89 |
| GCF_000703 125.1 | 0.69 | 0.4 | 0.0 | 25.13 | 38.94 | 10.13 |
| GCF_000526 495.1 | 0.34 | 0.4 | 0.0 | 20.02 | 20.63 | 13.67 |
| GCF_000389 635.1 | 0.39 | 3.0 | 0.0 | 21.05 | 0.02 | 13.71 |
| GCF_000307 125.1 | 0.00 | 0.0 | 0.0 | 21.66 | 0.11 | 10.89 |
| GCF_000284 435.1 | 0.00 | 0.8 | 0.0 | 21.61 | 40.35 | 4.79 |
| GCF_000283 555.1 | 0.99 | 0.4 | 0.0 | 21.58 | 0.10 | 4.73 |
| GCF_000270 | 0.00 | 0.8 | 0.0 | 21.59 | 40.51 | 6.8 |

| | | | | | | |
|---|---|---|---|---|---|---|
| 205.1 | | | | | | |
| GCF_000247605.1 | 1.06 | 0.8 | 0.0 | 22.95 | 0.01 | 11.82 |
| GCF_000244875.1 | 1.35 | 1.1 | 0.0 | 19.74 | 0.01 | 15.79 |
| GCF_000237085.1 | 0.00 | 0.4 | 0.0 | 18.76 | 0.03 | 15.83 |
| GCF_000218855.1 | 0.00 | 1.1 | 0.0 | 20.95 | 0.07 | 11.19 |
| GCF_000191905.1 | 0.00 | 1.1 | 0.0 | 21.07 | 0.06 | 11.19 |
| GCF_000190635.1 | 0.34 | 0.0 | 0.0 | 24.12 | 0.02 | 10.59 |
| GCF_000184705.1 | 0.00 | 0.0 | 0.0 | 21.68 | 0.12 | 11.25 |
| GCF_000166695.1 | 0.00 | 0.0 | 0.0 | 23.29 | 1.97 | 10.13 |
| GCF_000165465.1 | 0.87 | 0.8 | 0.0 | 29.02 | 0.12 | 9.3 |
| GCF_000022325.1 | 0.00 | 0.0 | 0.0 | 21.66 | 1.02 | 10.56 |
| GCF_000020165.1 | 0.00 | 0.4 | 0.0 | 21.26 | 0.12 | 10.89 |
| GCF_000019165.1 | 0.89 | 0.5 | 0.0 | 25.72 | 0.01 | 9.97 |
| GCF_000016165.1 | 1.27 | 1.5 | 0.0 | 25.90 | 0.00 | 10.59 |
| GCF_000014125.1 | 0.00 | 0.4 | 0.0 | 26.64 | 0.19 | 9.55 |
| GCF_000013845.2 | 0.00 | 0.4 | 0.0 | 23.20 | 0.26 | 9.76 |
| GCF_000012865.1 | 0.00 | 0.6 | 0.0 | 29.44 | 0.00 | 10.31 |
| GCF_000009685.1 | 0.00 | 0.4 | 0.0 | 23.25 | 0.35 | 10.13 |
| GCF_000008765.1 | 0.00 | 1.1 | 0.0 | 21.07 | 0.05 | 11.19 |
| GCF_000512 | 0.47 | 0.4 | 0.0 | 24.70 | 0.07 | 10.47 |

| | | | | | | |
|---|---|---|---|---|---|---|
| 895.1 | | | | | | |
| GCF_000305815.1 | 0.00 | 0.8 | 0.0 | 24.30 | 6.19 | 10.52 |
| GCF_000305775.1 | 0.00 | 0.8 | 0.0 | 23.79 | 7.12 | 10.52 |
| GCF_002005145.1 | 0.84 | 1.1 | 0.0 | 28.12 | 0.05 | 10.03 |
| GCF_900184125.1 | 0.00 | 0.0 | 0.0 | 21.47 | 1.12 | 0.03 |
| GCF_900166995.1 | 0.00 | 0.0 | 0.0 | 21.35 | 1.03 | 0.02 |
| GCF_002564135.1 | 0.00 | 0.0 | 0.0 | 21.47 | 1.08 | 10.56 |
| GCF_002300505.1 | 0.00 | 0.2 | 0.0 | 21.90 | 1.02 | 10.56 |
| GCF_002300425.1 | 0.00 | 0.0 | 0.0 | 21.78 | 1.12 | 10.56 |
| GCF_002300395.1 | 0.00 | 0.0 | 0.0 | 21.50 | 1.11 | 10.56 |
| GCF_002300375.1 | 0.00 | 0.0 | 0.0 | 21.47 | 1.05 | 10.56 |
| GCF_002300335.1 | 0.00 | 0.0 | 0.0 | 21.80 | 1.02 | 10.56 |
| GCF_002564185.1 | 0.00 | 0.0 | 0.0 | 22.22 | 1.01 | 10.56 |
| GCF_003047205.1 | 1.19 | 1.8 | 0.0 | 28.73 | 0.00 | 10.22 |
| GCF_006380995.1 | 0.55 | 1.5 | 0.0 | 22.10 | 0.93 | 11.19 |
| GCF_018884745.1 | 0.38 | 1.5 | 0.0 | 22.05 | 0.48 | 0.54 |
| GCF_006381115.1 | 1.06 | 2.7 | 0.0 | 22.21 | 0.48 | 11.19 |
| GCF_002082985.1 | 0.45 | 1.5 | 0.02 | 21.51 | 0.88 | 11.19 |
| GCF_900184185.1 | 0.00 | 0.0 | 0.0 | 21.71 | 1.02 | 0.02 |
| GCF_900184 | 0.00 | 0.0 | 0.0 | 21.63 | 1.00 | 0.02 |

| | | | | | | |
|---|---|---|---|---|---|---|
| 135.1 | | | | | | |
| GCF_900168285.1 | 0.48 | 0.0 | 0.0 | 21.29 | 1.05 | 0.02 |
| GCF_900168145.1 | 1.01 | 0.6 | 0.0 | 21.54 | 1.06 | 0.92 |
| GCF_900168085.1 | 0.24 | 0.0 | 0.0 | 21.75 | 1.08 | 0.02 |
| GCF_009905255.1 | 0.00 | 0.4 | 0.0 | 28.73 | 4.85 | 0.0 |
| GCF_000955725.1 | 0.00 | 0.0 | 0.0 | 22.63 | 62.21 | 10.51 |
| GCF_000806225.2 | 1.00 | 0.8 | 0.0 | 29.94 | 60.63 | 9.93 |
| GCF_000307585.2 | 0.00 | 0.4 | 0.0 | 27.37 | 6.91 | 10.13 |
| GCF_000189775.2 | 0.96 | 0.8 | 0.0 | 29.22 | 0.07 | 9.65 |
| GCF_000175295.2 | 0.24 | 0.4 | 0.0 | 31.79 | 5.69 | 9.83 |
| GCF_000166775.1 | 0.00 | 0.0 | 0.0 | 22.70 | 0.91 | 10.56 |
| GCF_000166355.1 | 0.00 | 0.0 | 0.0 | 23.93 | 0.46 | 10.13 |
| GCF_000166335.1 | 0.00 | 0.0 | 0.0 | 24.64 | 0.33 | 10.3 |
| GCF_000147695.2 | 0.00 | 0.6 | 0.0 | 29.52 | 1.54 | 9.93 |
| GCF_000145215.1 | 0.00 | 0.0 | 0.0 | 24.59 | 0.38 | 10.45 |
| GCF_000025645.1 | 0.00 | 0.0 | 0.0 | 30.71 | 1.55 | 9.95 |
| GCF_000019085.1 | 0.24 | 0.4 | 0.0 | 31.85 | 6.16 | 9.85 |
| GCF_000019065.1 | 0.00 | 0.6 | 0.0 | 32.72 | 45.11 | 9.94 |
| GCF_902386365.1 | 0.37 | 1.1 | 0.0 | 22.83 | 0.47 | 0.85 |
| GCF_900446 | 0.37 | 1.5 | 0.02 | 22.12 | 0.52 | 0.4 |

| | | | | | | |
|---|---|---|---|---|---|---|
| 965.1 | | | | | | |
| GCF_021378415.1 | 0.37 | 1.5 | 0.0 | 22.38 | 0.31 | 11.19 |
| GCF_019095765.1 | 0.45 | 1.5 | 0.0 | 22.50 | 0.38 | 0.76 |
| GCF_018885105.1 | 0.37 | 1.5 | 0.0 | 22.31 | 0.39 | 0.26 |
| GCF_018884965.1 | 0.53 | 1.5 | 0.0 | 23.38 | 0.42 | 0.54 |
| GCF_018884885.1 | 0.32 | 1.5 | 0.0 | 22.18 | 0.43 | 0.25 |
| GCF_018884765.1 | 0.37 | 1.5 | 0.02 | 22.31 | 0.59 | 0.46 |
| GCF_018255775.1 | 0.37 | 1.5 | 0.0 | 23.09 | 0.36 | 0.84 |
| GCF_015732555.1 | 0.37 | 1.5 | 0.0 | 22.76 | 0.43 | 1.18 |
| GCF_015238635.1 | 0.44 | 1.5 | 0.0 | 21.71 | 0.47 | 0.66 |
| GCF_014236775.1 | 0.37 | 1.5 | 0.0 | 22.52 | 0.53 | 11.19 |
| GCF_009867095.1 | 0.70 | 1.5 | 0.0 | 22.20 | 0.39 | 0.52 |
| GCF_009730495.1 | 0.37 | 1.5 | 0.0 | 22.73 | 0.44 | 11.19 |
| GCF_008245165.1 | 0.37 | 1.5 | 0.0 | 22.25 | 0.29 | 11.19 |
| GCF_006408335.1 | 0.62 | 2.3 | 0.0 | 23.86 | 1.27 | 11.19 |
| GCF_006408325.1 | 0.40 | 1.5 | 0.0 | 22.90 | 0.38 | 11.19 |
| GCF_006381095.1 | 0.37 | 1.5 | 0.0 | 22.82 | 0.46 | 11.19 |
| GCF_006381025.1 | 0.57 | 1.9 | 0.0 | 22.77 | 0.40 | 11.19 |
| GCF_006379355.1 | 0.37 | 1.5 | 0.0 | 22.55 | 0.45 | 11.19 |
| GCF_003482 | 0.37 | 1.5 | 0.0 | 23.50 | 0.32 | 11.19 |

| | | | | | | |
|---|---|---|---|---|---|---|
| 345.1 | | | | | | |
| GCF_003482 325.1 | 0.37 | 1.5 | 0.0 | 22.41 | 0.27 | 11.19 |
| GCF_003482 255.1 | 0.45 | 1.5 | 0.0 | 22.02 | 0.48 | 11.19 |
| GCF_003482 035.1 | 0.37 | 1.5 | 0.0 | 22.51 | 0.35 | 11.19 |
| GCF_003481 965.1 | 0.32 | 1.1 | 0.0 | 22.35 | 0.47 | 11.19 |
| GCF_003481 905.1 | 0.37 | 1.5 | 0.0 | 21.92 | 0.56 | 11.19 |
| GCF_003313 585.1 | 0.37 | 1.5 | 0.0 | 23.01 | 0.40 | 11.19 |
| GCF_003313 565.1 | 0.37 | 1.5 | 0.0 | 23.21 | 0.47 | 11.19 |
| GCF_003313 545.1 | 0.37 | 1.5 | 0.0 | 22.79 | 0.37 | 11.19 |
| GCF_002946 195.1 | 0.37 | 1.5 | 0.0 | 22.22 | 0.30 | 11.19 |
| GCF_002946 135.1 | 0.37 | 1.5 | 0.0 | 22.41 | 0.38 | 11.19 |
| GCF_002945 945.1 | 0.37 | 1.5 | 0.0 | 22.29 | 0.35 | 11.19 |
| GCF_002945 755.1 | 0.37 | 1.5 | 0.0 | 22.91 | 0.38 | 11.19 |
| GCF_002945 665.1 | 0.37 | 1.5 | 0.0 | 22.68 | 0.29 | 11.19 |
| GCF_002945 515.1 | 0.37 | 1.5 | 0.0 | 22.86 | 0.43 | 11.19 |
| GCF_002945 415.1 | 0.37 | 1.5 | 0.0 | 22.33 | 0.38 | 11.19 |
| GCF_002812 585.1 | 0.37 | 1.1 | 0.0 | 22.83 | 0.47 | 11.19 |
| GCF_002080 065.1 | 0.37 | 1.5 | 0.0 | 23.35 | 0.47 | 11.19 |
| GCF_001972 015.1 | 0.37 | 1.5 | 0.0 | 22.63 | 0.42 | 11.19 |
| GCF_000953 | 0.37 | 1.5 | 0.0 | 23.45 | 0.45 | 11.19 |

| | | | | | | |
|---|---|---|---|---|---|---|
| 275.1 | | | | | | |
| GCF_000932055.2 | 0.37 | 1.5 | 0.0 | 23.08 | 0.49 | 11.19 |
| GCF_000826625.1 | 0.37 | 1.5 | 0.0 | 22.50 | 0.45 | 11.19 |
| GCF_000085225.1 | 0.37 | 1.5 | 0.0 | 22.62 | 0.30 | 11.19 |
| GCF_000009205.2 | 0.37 | 1.5 | 0.0 | 23.28 | 0.44 | 11.19 |
| GCF_003999255.1 | 0.16 | 0.2 | 0.0 | 23.43 | 0.50 | 10.56 |
| GCF_000016545.1 | 0.00 | 0.0 | 0.0 | 23.61 | 0.66 | 10.56 |
| GCF_003490105.1 | 0.71 | 1.5 | 0.0 | 21.79 | 0.36 | 11.19 |
| GCF_003482065.1 | 0.91 | 1.5 | 0.0 | 23.24 | 0.43 | 11.19 |
| GCF_001972105.1 | 0.38 | 1.5 | 0.0 | 21.91 | 0.79 | 11.19 |
| GCF_000092945.1 | 1.28 | 1.1 | 0.0 | 26.19 | 0.01 | 10.16 |
| GCF_014170115.1 | 0.23 | 0.4 | 0.0 | 27.57 | 0.06 | 0.16 |
| GCF_001679705.1 | 0.23 | 0.4 | 0.0 | 27.39 | 0.04 | 10.54 |
| GCF_001642655.1 | 0.23 | 0.4 | 0.0 | 26.94 | 0.04 | 10.54 |
| GCF_006408435.1 | 0.64 | 1.5 | 0.0 | 22.65 | 0.44 | 11.19 |
| GCF_003482225.1 | 0.84 | 1.5 | 0.0 | 21.82 | 0.41 | 11.19 |
| GCF_018885065.1 | 0.38 | 1.5 | 0.0 | 22.13 | 0.54 | 11.19 |
| GCF_018885085.1 | 0.53 | 1.5 | 0.0 | 21.78 | 0.37 | 0.21 |
| GCF_006408375.1 | 0.53 | 1.5 | 0.02 | 21.84 | 0.38 | 11.19 |
| GCF_002812 | 0.55 | 1.1 | 0.0 | 22.41 | 0.31 | 11.19 |

| | | | | | | |
|---|---|---|---|---|---|---|
| 645.1 | | | | | | |
| GCF_013487845.1 | 0.00 | 0.5 | 0.0 | 16.86 | 0.02 | 9.09 |
| GCF_018884825.1 | 0.55 | 1.5 | 0.0 | 22.65 | 0.24 | 0.16 |
| GCF_006408285.1 | 0.55 | 1.5 | 0.0 | 22.62 | 1.72 | 11.19 |
| GCF_002812605.1 | 0.55 | 1.1 | 0.0 | 22.85 | 0.35 | 11.19 |
| GCF_002007885.1 | 0.55 | 1.5 | 0.0 | 22.40 | 0.33 | 11.19 |
| GCF_001971945.1 | 0.55 | 1.5 | 0.0 | 22.51 | 0.32 | 11.19 |
| GCF_002945855.1 | 0.37 | 1.5 | 0.0 | 22.43 | 0.36 | 11.19 |
| GCF_902387955.1 | 0.24 | 0.0 | 0.0 | 23.38 | 0.20 | 12.21 |
| GCF_000225345.1 | 0.24 | 0.0 | 0.0 | 23.38 | 0.20 | 12.21 |
| GCF_009796305.1 | 0.00 | 0.4 | 0.0 | 24.39 | 0.03 | 10.52 |
| GCF_900604355.1 | 0.23 | 0.0 | 0.0 | 23.70 | 21.01 | 4.29 |
| GCF_020181455.1 | 0.39 | 0.4 | 0.0 | 23.88 | 0.49 | 0.97 |
| GCF_016904155.1 | 0.00 | 0.4 | 0.0 | 25.61 | 0.30 | 10.66 |
| GCF_016904015.1 | 0.00 | 0.0 | 0.0 | 27.02 | 0.38 | 10.61 |
| GCF_016889005.1 | 0.00 | 0.0 | 0.0 | 25.26 | 0.04 | 1.41 |
| GCF_009831375.1 | 0.00 | 0.0 | 0.0 | 25.27 | 0.37 | 10.66 |
| GCF_009684695.1 | 0.58 | 0.0 | 0.0 | 23.57 | 0.08 | 12.21 |
| GCF_004295125.1 | 1.17 | 1.1 | 0.0 | 25.27 | 0.04 | 12.21 |
| GCF_002234 | 0.55 | 1.5 | 0.0 | 22.49 | 0.32 | 11.19 |

| | | | | | | |
|---|---|---|---|---|---|---|
| 355.1 | | | | | | |
| GCF_011058 795.1 | 0.44 | 0.0 | 0.0 | 26.43 | 0.13 | 3.19 |
| GCF_011058 775.1 | 0.44 | 0.0 | 0.0 | 26.83 | 0.06 | 3.19 |
| GCF_011058 755.1 | 0.44 | 0.2 | 0.0 | 25.13 | 0.32 | 0.1 |
| GCF_011058 735.1 | 0.44 | 0.2 | 0.0 | 25.84 | 0.15 | 5.08 |
| GCF_011058 715.1 | 0.44 | 0.0 | 0.0 | 26.36 | 0.16 | 3.28 |
| GCF_009857 205.1 | 0.44 | 0.0 | 0.0 | 26.45 | 1.24 | 3.21 |
| GCF_009649 955.1 | 1.83 | 0.5 | 0.0 | 23.27 | 0.00 | 10.51 |
| GCF_002250 075.1 | 1.99 | 1.5 | 0.0 | 27.15 | 0.07 | 10.13 |
| GCF_000145 615.1 | 0.48 | 0.2 | 0.0 | 27.95 | 0.09 | 10.13 |
| GCF_001972 085.1 | 0.55 | 1.5 | 0.0 | 22.27 | 0.29 | 11.19 |
| GCF_006454 545.1 | 0.37 | 1.5 | 0.0 | 22.94 | 0.45 | 11.19 |
| GCF_002946 035.1 | 0.37 | 1.5 | 0.0 | 22.34 | 0.33 | 11.19 |
| GCF_009903 525.1 | 0.42 | 1.5 | 0.0 | 22.05 | 0.39 | 11.19 |
| GCF_006380 905.1 | 0.42 | 1.5 | 0.0 | 21.97 | 0.39 | 11.19 |
| GCF_006379 335.1 | 0.42 | 1.5 | 0.0 | 21.89 | 0.43 | 11.19 |
| GCF_902387 945.1 | 0.00 | 0.0 | 0.0 | 25.51 | 0.06 | 10.13 |
| GCF_900461 315.1 | 0.63 | 0.0 | 0.0 | 23.81 | 0.00 | 15.98 |
| GCF_900461 275.1 | 0.00 | 0.0 | 0.0 | 20.68 | 18.47 | 1.55 |
| GCF_900105 | 0.63 | 0.4 | 0.0 | 22.16 | 3.69 | 0.65 |

| | | | | | | |
|---|---|---|---|---|---|---|
| 215.1 | | | | | | |
| GCF_020735365.1 | 0.32 | 0.4 | 0.0 | 23.62 | 1.88 | 1.41 |
| GCF_018866245.1 | 0.00 | 0.4 | 0.0 | 26.80 | 0.01 | 0.7 |
| GCF_005848555.1 | 0.32 | 0.4 | 0.0 | 24.59 | 0.58 | 12.21 |
| GCF_003589745.1 | 0.00 | 0.0 | 0.0 | 25.51 | 0.06 | 10.13 |
| GCF_002797975.1 | 0.00 | 0.0 | 0.0 | 24.07 | 5.58 | 15.98 |
| GCF_000686105.1 | 0.95 | 1.1 | 0.0 | 16.92 | 17.19 | 23.71 |
| GCF_000526995.1 | 0.95 | 0.0 | 0.0 | 21.98 | 30.77 | 17.58 |
| GCF_000214435.1 | 0.00 | 1.1 | 0.0 | 29.05 | 0.00 | 9.85 |
| GCF_000159975.2 | 0.00 | 0.0 | 0.0 | 22.20 | 29.11 | 13.24 |
| GCF_000144625.1 | 0.00 | 0.4 | 0.0 | 23.52 | 0.00 | 15.63 |
| GCF_900461125.1 | 0.00 | 0.0 | 0.0 | 20.77 | 5.63 | 17.58 |
| GCF_020215665.1 | 0.00 | 0.0 | 0.0 | 20.09 | 0.15 | 3.89 |
| GCF_014131715.1 | 0.00 | 0.0 | 0.0 | 20.14 | 0.05 | 4.32 |
| GCF_013112015.1 | 1.91 | 0.0 | 0.0 | 20.24 | 0.03 | 1.05 |
| GCF_010669205.1 | 0.00 | 0.0 | 0.0 | 20.25 | 0.04 | 4.33 |
| GCF_004210255.1 | 0.21 | 0.0 | 0.0 | 19.83 | 0.04 | 17.58 |
| GCF_002222595.2 | 0.00 | 0.0 | 0.0 | 24.02 | 0.76 | 10.61 |
| GCF_001689125.2 | 1.91 | 0.0 | 0.0 | 20.89 | 0.02 | 15.77 |
| GCF_000231 | 0.98 | 0.0 | 0.0 | 25.24 | 0.00 | 10.47 |

| | | | | | | |
|---|---|---|---|---|---|---|
| 405.2 | | | | | | |
| GCF_001577 795.1 | 0.38 | 1.5 | 0.0 | 22.46 | 0.41 | 11.19 |
| GCF_000014 405.1 | 0.00 | 0.5 | 0.0 | 4.34 | 0.03 | 9.15 |
| GCF_005886 215.1 | 0.95 | 0.0 | 0.0 | 18.61 | 0.00 | 16.01 |
| GCF_003482 125.1 | 0.76 | 1.5 | 0.0 | 21.92 | 0.44 | 11.19 |
| GCF_902388 025.1 | 1.34 | 0.4 | 0.0 | 24.81 | 0.35 | 1.04 |
| GCF_900215 405.1 | 0.00 | 0.0 | 0.0 | 21.47 | 0.15 | 10.63 |
| GCF_020735 705.1 | 0.00 | 0.8 | 0.0 | 21.33 | 0.12 | 1.59 |
| GCF_020181 555.1 | 0.81 | 0.4 | 0.0 | 19.73 | 0.04 | 15.7 |
| GCF_010508 875.1 | 0.13 | 0.4 | 0.0 | 21.28 | 0.03 | 15.39 |
| GCF_008727 735.1 | 0.00 | 0.4 | 0.0 | 24.76 | 0.02 | 10.09 |
| GCF_005473 905.2 | 0.34 | 0.0 | 0.0 | 18.48 | 0.03 | 13.67 |
| GCF_002564 085.1 | 0.00 | 0.0 | 0.0 | 21.59 | 0.13 | 10.58 |
| GCF_001998 765.1 | 1.34 | 0.4 | 0.0 | 24.81 | 0.35 | 10.61 |
| GCF_001692 755.1 | 0.00 | 0.0 | 0.0 | 21.55 | 0.15 | 10.58 |
| GCF_001688 625.2 | 0.13 | 0.4 | 0.0 | 22.25 | 0.03 | 15.31 |
| GCF_000255 615.2 | 0.00 | 0.0 | 0.0 | 21.59 | 0.13 | 10.58 |
| GCF_000184 925.1 | 0.00 | 0.0 | 0.0 | 21.54 | 0.11 | 10.58 |
| GCF_000015 865.1 | 0.67 | 0.0 | 0.0 | 21.03 | 0.14 | 10.98 |
| GCF_000022 | 1.01 | 0.8 | 0.0 | 21.22 | 0.01 | 11.82 |

| | | | | | | |
|---|---|---|---|---|---|---|
| 065.1 | | | | | | |
| GCF_020297 485.1 | 1.69 | 1.9 | 0.0 | 22.38 | 48.00 | 4.78 |
| GCF_016889 665.1 | 0.00 | 0.4 | 0.0 | 21.31 | 0.07 | 4.41 |
| GCF_013112 035.1 | 0.00 | 0.0 | 0.0 | 20.74 | 0.05 | 20.37 |
| GCF_006538 465.1 | 0.00 | 1.1 | 0.0 | 21.47 | 4.71 | 17.33 |
| GCF_002234 575.2 | 0.00 | 0.4 | 0.0 | 21.20 | 0.08 | 22.12 |
| GCF_001688 665.2 | 1.27 | 1.9 | 0.0 | 19.15 | 1.90 | 22.12 |
| GCF_000210 455.1 | 0.37 | 1.5 | 0.0 | 23.18 | 0.39 | 11.19 |
| GCF_020731 185.1 | 0.00 | 0.8 | 0.0 | 16.98 | 10.30 | 2.29 |
| GCF_006454 575.1 | 0.74 | 2.3 | 0.0 | 22.00 | 0.36 | 11.19 |
| GCF_021495 975.1 | 1.38 | 1.5 | 0.0 | 23.20 | 32.31 | 1.37 |
| GCF_020099 295.1 | 1.38 | 1.5 | 0.0 | 22.86 | 34.66 | 0.58 |
| GCF_016838 665.1 | 0.69 | 0.8 | 0.0 | 22.00 | 0.61 | 0.73 |
| GCF_016798 345.1 | 0.00 | 1.1 | 0.0 | 23.42 | 0.56 | 0.81 |
| GCF_014068 615.1 | 0.00 | 1.1 | 0.0 | 22.96 | 0.57 | 0.66 |
| GCF_009733 885.1 | 0.00 | 1.1 | 0.0 | 23.25 | 0.56 | 10.7 |
| GCF_003345 335.1 | 0.13 | 0.8 | 0.0 | 21.42 | 0.61 | 11.19 |
| GCF_003345 315.1 | 0.69 | 1.1 | 0.0 | 22.64 | 0.54 | 11.19 |
| GCF_003058 445.1 | 0.00 | 1.1 | 0.0 | 22.98 | 0.55 | 10.7 |
| GCF_003058 | 0.00 | 1.1 | 0.0 | 23.36 | 0.50 | 10.7 |

| | | | | | | |
|---|---|---|---|---|---|---|
| 345.1 | | | | | | |
| GCF_003017225.1 | 0.00 | 1.1 | 0.0 | 22.82 | 3.89 | 11.19 |
| GCF_003017195.1 | 0.00 | 0.8 | 0.0 | 23.34 | 4.61 | 11.19 |
| GCF_003017145.1 | 0.00 | 0.8 | 0.0 | 23.53 | 4.43 | 11.19 |
| GCF_002866225.1 | 1.38 | 1.1 | 0.0 | 22.12 | 0.61 | 11.19 |
| GCF_002866125.1 | 0.69 | 1.1 | 0.0 | 22.31 | 0.52 | 11.19 |
| GCF_002866045.1 | 0.08 | 1.1 | 0.0 | 21.72 | 0.56 | 11.19 |
| GCF_002865885.1 | 0.00 | 1.1 | 0.0 | 22.07 | 0.54 | 11.19 |
| GCF_002865845.1 | 0.00 | 1.1 | 0.0 | 22.28 | 0.57 | 11.19 |
| GCF_002865805.1 | 0.69 | 1.1 | 0.0 | 22.71 | 0.49 | 10.7 |
| GCF_002865785.1 | 0.08 | 1.1 | 0.0 | 22.04 | 28.40 | 11.19 |
| GCF_002865765.1 | 0.00 | 1.1 | 0.0 | 22.90 | 0.51 | 10.7 |
| GCF_002865745.1 | 0.00 | 1.1 | 0.0 | 22.72 | 0.52 | 10.7 |
| GCF_001921965.1 | 0.00 | 1.1 | 0.0 | 22.56 | 0.60 | 11.19 |
| GCF_001921945.1 | 0.77 | 1.1 | 0.0 | 21.70 | 0.60 | 11.19 |
| GCF_001921905.1 | 0.69 | 1.1 | 0.0 | 22.84 | 0.61 | 10.98 |
| GCF_001889345.1 | 0.00 | 1.1 | 0.0 | 23.13 | 0.58 | 10.7 |
| GCF_001886775.1 | 0.69 | 1.1 | 0.0 | 21.63 | 8.16 | 11.19 |
| GCF_001879625.1 | 0.69 | 1.1 | 0.0 | 22.80 | 0.59 | 11.19 |
| GCF_001879 | 0.00 | 1.1 | 0.0 | 22.31 | 0.57 | 11.19 |

| | | | | | | |
|---|---|---|---|---|---|---|
| 605.1 | | | | | | |
| GCF_000829 015.1 | 0.00 | 1.1 | 0.0 | 22.97 | 0.57 | 10.7 |
| GCF_000817 935.1 | 0.08 | 1.1 | 0.0 | 20.62 | 0.60 | 11.19 |
| GCF_000063 585.1 | 0.00 | 1.1 | 0.0 | 23.05 | 0.54 | 10.7 |
| GCF_000022 765.1 | 0.00 | 1.1 | 0.0 | 22.13 | 0.58 | 11.19 |
| GCF_000020 345.1 | 0.05 | 1.1 | 0.0 | 21.36 | 0.54 | 11.19 |
| GCF_000019 545.1 | 0.08 | 1.1 | 0.0 | 21.49 | 0.45 | 11.19 |
| GCF_000019 305.1 | 0.00 | 1.1 | 0.0 | 22.06 | 0.57 | 11.19 |
| GCF_000017 045.1 | 0.00 | 1.1 | 0.0 | 23.73 | 0.49 | 10.7 |
| GCF_000017 025.1 | 0.00 | 1.1 | 0.0 | 23.36 | 0.52 | 10.7 |
| GCF_000016 505.1 | 0.00 | 0.4 | 0.0 | 22.00 | 0.12 | 10.98 |
| GCF_000010 265.1 | 0.00 | 0.4 | 0.0 | 22.50 | 0.11 | 10.98 |
| GCF_020641 055.1 | 0.00 | 1.5 | 0.0 | 20.97 | 25.55 | 0.14 |
| GCF_002441 855.2 | 0.00 | 0.8 | 0.0 | 21.79 | 1.07 | 13.44 |
| GCF_001481 725.1 | 0.00 | 0.8 | 0.0 | 21.78 | 0.47 | 13.45 |
| GCF_000807 675.2 | 0.00 | 0.8 | 0.0 | 21.63 | 0.06 | 13.44 |
| GCF_000152 245.2 | 0.00 | 0.8 | 0.0 | 21.45 | 0.45 | 13.44 |
| GCF_019884 785.1 | 1.77 | 1.5 | 0.0 | 23.77 | 0.03 | 3.67 |
| GCF_019203 185.1 | 1.06 | 0.8 | 0.0 | 26.97 | 0.07 | 0.09 |
| GCF_004103 | 0.35 | 0.4 | 0.0 | 20.86 | 0.01 | 10.18 |

| | | | | | | |
|---|---|---|---|---|---|---|
| 735.1 | | | | | | |
| GCF_002998925.1 | 0.00 | 0.4 | 0.0 | 5.91 | 0.10 | 7.37 |
| GCF_002080475.1 | 0.71 | 0.8 | 0.0 | 23.45 | 0.03 | 12.04 |
| GCF_001854185.1 | 0.71 | 0.8 | 0.0 | 23.28 | 0.03 | 12.04 |
| GCF_001042715.1 | 0.71 | 1.1 | 0.0 | 24.01 | 0.02 | 11.74 |
| GCF_000024205.1 | 0.16 | 1.9 | 0.0 | 19.22 | 0.00 | 15.54 |
| GCF_006228565.1 | 0.00 | 1.3 | 0.0 | 27.60 | 0.00 | 10.2 |
| GCF_001729945.1 | 1.83 | 2.1 | 0.0 | 25.40 | 0.00 | 9.83 |
| GCF_001267405.1 | 0.00 | 1.3 | 0.0 | 28.23 | 0.00 | 10.19 |
| GCF_000231385.2 | 0.51 | 1.5 | 0.0 | 22.56 | 0.01 | 13.94 |
| GCF_900636895.1 | 0.00 | 0.5 | 0.0 | 27.48 | 3.04 | 9.29 |
| GCF_020995385.1 | 0.15 | 0.5 | 0.0 | 26.69 | 49.39 | 0.04 |
| GCF_020991185.1 | 0.00 | 0.5 | 0.0 | 27.64 | 0.55 | 0.01 |
| GCF_017840575.1 | 0.00 | 0.5 | 0.0 | 25.36 | 15.72 | 0.55 |
| GCF_017638885.1 | 0.78 | 0.5 | 0.0 | 25.40 | 0.43 | 0.11 |
| GCF_017498665.1 | 0.00 | 0.5 | 0.0 | 27.68 | 2.21 | 0.09 |
| GCF_014841035.1 | 0.39 | 0.5 | 0.0 | 26.34 | 0.19 | 9.2 |
| GCF_009834385.1 | 0.00 | 0.5 | 0.0 | 26.07 | 11.92 | 9.22 |
| GCF_007876425.1 | 1.29 | 1.2 | 0.0 | 25.79 | 3.78 | 9.22 |
| GCF_005886 | 0.00 | 0.5 | 0.0 | 26.10 | 2.91 | 9.32 |

| | | | | | | |
|---|---|---|---|---|---|---|
| 075.1 | | | | | | |
| GCF_003584685.1 | 0.00 | 0.5 | 0.0 | 26.09 | 3.12 | 9.16 |
| GCF_003428395.1 | 0.00 | 0.2 | 0.0 | 27.14 | 0.09 | 9.15 |
| GCF_003316915.1 | 0.00 | 0.5 | 0.0 | 27.97 | 0.10 | 9.15 |
| GCF_002176855.1 | 0.00 | 0.5 | 0.0 | 26.40 | 0.24 | 9.29 |
| GCF_002176835.1 | 0.78 | 0.7 | 0.0 | 25.81 | 0.36 | 9.19 |
| GCF_000439915.2 | 0.00 | 0.5 | 0.0 | 28.35 | 0.85 | 9.15 |
| GCF_000204985.1 | 0.00 | 0.5 | 0.0 | 27.44 | 0.17 | 9.19 |
| GCF_000091405.1 | 0.00 | 0.2 | 0.0 | 27.93 | 0.17 | 7.38 |
| GCF_000008065.1 | 0.00 | 0.5 | 0.0 | 26.17 | 0.24 | 9.32 |
| GCF_902459515.1 | 0.00 | 0.4 | 0.0 | 22.81 | 0.44 | 10.13 |
| GCF_900604345.1 | 0.81 | 0.4 | 0.0 | 24.58 | 0.02 | 10.13 |
| GCF_900475545.1 | 0.09 | 0.8 | 0.0 | 22.90 | 0.52 | 0.05 |
| GCF_900465025.1 | 1.61 | 1.1 | 0.0 | 22.58 | 3.71 | 2.36 |
| GCF_900461205.1 | 0.00 | 0.8 | 0.0 | 22.34 | 0.45 | 0.12 |
| GCF_900168365.1 | 0.00 | 0.4 | 0.0 | 24.45 | 0.25 | 0.09 |
| GCF_900010805.1 | 1.61 | 2.3 | 0.0 | 19.67 | 0.15 | 0.87 |
| GCF_020911725.1 | 1.61 | 0.4 | 0.0 | 24.72 | 0.04 | 10.98 |
| GCF_020139275.1 | 0.00 | 0.8 | 0.0 | 21.81 | 0.59 | 0.06 |
| GCF_018223 | 0.81 | 1.9 | 0.0 | 20.48 | 1.14 | 13.78 |

| | | | | | | |
|---|---|---|---|---|---|---|
| 745.1 | | | | | | |
| GCF_018140655.1 | 0.00 | 1.1 | 0.0 | 22.51 | 0.09 | 13.82 |
| GCF_018140595.1 | 1.61 | 2.3 | 0.0 | 20.54 | 0.60 | 0.52 |
| GCF_016027375.1 | 0.00 | 0.8 | 0.0 | 22.42 | 0.49 | 0.12 |
| GCF_016026995.1 | 0.00 | 0.4 | 0.0 | 22.43 | 0.60 | 0.11 |
| GCF_015643605.1 | 1.61 | 2.3 | 0.0 | 20.19 | 0.65 | 15.9 |
| GCF_014137775.1 | 0.81 | 0.8 | 0.0 | 19.32 | 0.08 | 0.05 |
| GCF_014137745.1 | 0.00 | 1.1 | 0.0 | 22.08 | 0.10 | 1.13 |
| GCF_014137665.1 | 0.81 | 0.4 | 0.0 | 19.31 | 0.07 | 0.06 |
| GCF_014131795.1 | 0.27 | 1.1 | 0.0 | 21.73 | 0.07 | 1.81 |
| GCF_013407635.1 | 1.75 | 2.3 | 0.0 | 20.23 | 0.68 | 0.62 |
| GCF_013407215.1 | 1.69 | 2.3 | 0.0 | 20.71 | 0.52 | 0.25 |
| GCF_013297265.1 | 1.69 | 2.3 | 0.0 | 20.57 | 0.51 | 15.9 |
| GCF_013297175.1 | 1.69 | 2.3 | 0.0 | 20.81 | 0.53 | 0.24 |
| GCF_013297125.1 | 1.69 | 1.9 | 0.0 | 19.18 | 0.11 | 0.93 |
| GCF_013297065.1 | 1.69 | 2.3 | 0.0 | 20.61 | 0.49 | 15.9 |
| GCF_013296465.1 | 1.69 | 2.3 | 0.0 | 20.64 | 0.48 | 0.26 |
| GCF_013296445.1 | 1.69 | 2.3 | 0.0 | 20.61 | 0.57 | 15.9 |
| GCF_013296425.1 | 1.69 | 2.3 | 0.0 | 20.79 | 0.52 | 15.9 |
| GCF_013296 | 1.61 | 2.3 | 0.0 | 19.59 | 2.85 | 0.99 |

| | | | | | | |
|---|---|---|---|---|---|---|
| 395.1 | | | | | | |
| GCF_013294 755.1 | 1.69 | 2.3 | 0.0 | 20.63 | 0.52 | 15.9 |
| GCF_013294 745.1 | 1.69 | 2.3 | 0.0 | 20.59 | 0.47 | 0.23 |
| GCF_013294 705.1 | 1.69 | 2.3 | 0.0 | 20.76 | 0.52 | 0.22 |
| GCF_013294 675.1 | 1.69 | 2.3 | 0.0 | 20.60 | 0.51 | 15.9 |
| GCF_013294 665.1 | 1.69 | 2.3 | 0.0 | 20.61 | 0.51 | 0.23 |
| GCF_013294 605.1 | 1.69 | 2.3 | 0.0 | 20.71 | 0.49 | 0.24 |
| GCF_013294 555.1 | 1.61 | 2.3 | 0.0 | 19.61 | 2.02 | 0.72 |
| GCF_013294 515.1 | 1.69 | 2.3 | 0.0 | 20.46 | 0.48 | 15.9 |
| GCF_013294 505.1 | 1.69 | 2.3 | 0.0 | 20.58 | 0.49 | 15.9 |
| GCF_013294 475.1 | 1.69 | 2.3 | 0.0 | 20.79 | 0.58 | 0.2 |
| GCF_013294 465.1 | 1.69 | 2.3 | 0.0 | 20.59 | 0.48 | 0.23 |
| GCF_013294 445.1 | 1.69 | 1.9 | 0.0 | 19.03 | 0.12 | 0.87 |
| GCF_013294 405.1 | 1.61 | 2.3 | 0.0 | 20.46 | 5.70 | 0.36 |
| GCF_013294 375.1 | 1.61 | 2.3 | 0.0 | 19.50 | 0.15 | 0.88 |
| GCF_013294 345.1 | 1.61 | 2.3 | 0.0 | 19.56 | 2.94 | 1.03 |
| GCF_013294 325.1 | 1.69 | 2.3 | 0.0 | 20.68 | 0.51 | 15.9 |
| GCF_013294 205.1 | 1.61 | 2.3 | 0.0 | 19.51 | 0.17 | 15.9 |
| GCF_013294 145.1 | 1.69 | 2.3 | 0.0 | 20.68 | 0.47 | 0.23 |
| GCF_013149 | 0.81 | 0.8 | 0.0 | 19.42 | 0.07 | 0.04 |

| | | | | | | |
|---|---|---|---|---|---|---|
| 715.1 | | | | | | |
| GCF_013149705.1 | 0.81 | 0.8 | 0.0 | 19.36 | 0.07 | 0.06 |
| GCF_013149665.1 | 0.81 | 0.8 | 0.0 | 19.27 | 0.07 | 0.12 |
| GCF_013149605.1 | 0.81 | 0.8 | 0.0 | 19.26 | 0.07 | 0.07 |
| GCF_013149565.1 | 1.61 | 2.3 | 0.0 | 19.88 | 10.34 | 1.1 |
| GCF_013149465.1 | 0.81 | 0.8 | 0.0 | 19.50 | 0.07 | 13.67 |
| GCF_013149325.1 | 0.81 | 0.8 | 0.0 | 19.39 | 0.12 | 0.02 |
| GCF_013112415.1 | 0.00 | 1.1 | 0.0 | 22.18 | 0.14 | 1.13 |
| GCF_009650335.1 | 0.00 | 1.1 | 0.0 | 22.29 | 0.06 | 13.67 |
| GCF_009650315.1 | 0.00 | 1.1 | 0.0 | 22.16 | 0.06 | 13.67 |
| GCF_006742065.1 | 0.27 | 1.1 | 0.0 | 21.95 | 0.10 | 13.67 |
| GCF_006016075.1 | 0.81 | 0.8 | 0.0 | 29.01 | 0.06 | 10.13 |
| GCF_005145085.1 | 0.00 | 1.1 | 0.0 | 22.06 | 0.10 | 13.82 |
| GCF_004328885.1 | 0.00 | 0.4 | 0.0 | 24.40 | 0.28 | 10.13 |
| GCF_004101825.1 | 0.00 | 0.8 | 0.0 | 21.21 | 0.24 | 10.18 |
| GCF_003606265.1 | 0.16 | 0.8 | 0.0 | 21.40 | 0.20 | 10.18 |
| GCF_003315755.1 | 0.00 | 1.1 | 0.0 | 22.30 | 0.07 | 13.82 |
| GCF_002327205.1 | 0.00 | 0.4 | 0.0 | 24.05 | 0.26 | 10.13 |
| GCF_002327185.1 | 0.00 | 0.4 | 0.0 | 24.19 | 0.17 | 10.13 |
| GCF_002003 | 0.81 | 0.8 | 0.0 | 19.30 | 0.06 | 13.67 |

| | | | | | | |
|---|---|---|---|---|---|---|
| 385.1 | | | | | | |
| GCF_002003 365.1 | 0.81 | 0.8 | 0.0 | 19.46 | 0.08 | 13.67 |
| GCF_002003 345.1 | 1.69 | 2.3 | 0.0 | 20.85 | 0.49 | 15.9 |
| GCF_002003 325.1 | 0.81 | 0.8 | 0.0 | 19.29 | 0.07 | 13.67 |
| GCF_002003 305.1 | 1.29 | 1.9 | 0.0 | 19.84 | 0.04 | 15.98 |
| GCF_002003 285.1 | 0.81 | 0.8 | 0.0 | 19.39 | 0.07 | 13.67 |
| GCF_001991 075.2 | 1.61 | 0.8 | 0.0 | 24.88 | 0.10 | 10.13 |
| GCF_001886 875.1 | 0.00 | 1.1 | 0.0 | 22.13 | 0.17 | 13.82 |
| GCF_001646 605.1 | 0.00 | 1.1 | 0.0 | 22.36 | 0.08 | 13.82 |
| GCF_001456 065.2 | 0.00 | 1.5 | 0.0 | 22.12 | 0.10 | 13.82 |
| GCF_001304 735.1 | 0.44 | 2.3 | 0.0 | 22.45 | 0.54 | 10.13 |
| GCF_000878 275.1 | 0.00 | 1.1 | 0.0 | 22.27 | 0.15 | 13.82 |
| GCF_000833 105.2 | 1.61 | 2.3 | 0.0 | 20.79 | 0.42 | 15.98 |
| GCF_000827 955.1 | 0.00 | 0.4 | 0.0 | 23.23 | 0.13 | 10.59 |
| GCF_000827 935.1 | 0.00 | 0.4 | 0.0 | 23.11 | 0.13 | 10.59 |
| GCF_000767 745.1 | 1.61 | 2.3 | 0.0 | 20.35 | 0.60 | 13.78 |
| GCF_000473 995.1 | 0.81 | 0.8 | 0.0 | 19.30 | 0.07 | 13.67 |
| GCF_000340 885.1 | 1.69 | 1.9 | 0.0 | 19.05 | 0.11 | 16.01 |
| GCF_000020 285.1 | 0.00 | 0.4 | 0.0 | 22.76 | 0.13 | 10.7 |
| GCF_000016 | 1.61 | 2.3 | 0.0 | 20.46 | 0.62 | 13.78 |

| | | | | | | |
|---|---|---|---|---|---|---|
| 965.1 | | | | | | |
| GCF_000013285.1 | 0.00 | 0.8 | 0.0 | 22.66 | 0.54 | 10.13 |
| GCF_002812625.1 | 0.38 | 1.1 | 0.0 | 22.21 | 0.42 | 11.19 |
| GCF_018884665.1 | 0.37 | 1.5 | 0.02 | 22.00 | 0.49 | 0.48 |
| GCF_002082945.2 | 0.96 | 2.7 | 0.0 | 23.28 | 0.49 | 11.19 |
| GCF_000235605.1 | 0.51 | 2.3 | 0.0 | 22.97 | 0.00 | 15.98 |
| GCF_013781885.1 | 1.40 | 1.9 | 0.0 | 22.07 | 0.05 | 1.96 |
| GCF_001465175.1 | 0.00 | 1.1 | 0.0 | 22.42 | 0.09 | 13.82 |
| GCF_018884605.1 | 0.38 | 1.5 | 0.0 | 22.29 | 0.45 | 11.19 |
| GCF_000687955.1 | 0.70 | 1.1 | 0.0 | 23.30 | 33.06 | 10.13 |
| GCF_003482305.1 | 0.47 | 1.5 | 0.0 | 22.48 | 0.36 | 11.19 |
| GCF_017751145.1 | 1.75 | 1.2 | 0.0 | 24.39 | 0.65 | 1.15 |
| GCF_000020485.1 | 0.88 | 2.0 | 0.0 | 26.70 | 0.05 | 9.53 |
| GCF_013295945.1 | 1.69 | 1.9 | 0.0 | 21.00 | 0.55 | 15.9 |
| GCF_009903465.1 | 0.53 | 1.5 | 0.0 | 21.94 | 0.78 | 11.19 |
| GCF_010586925.1 | 0.78 | 1.0 | 0.0 | 24.95 | 0.21 | 9.26 |
| GCF_006380915.1 | 0.42 | 1.5 | 0.02 | 21.77 | 0.44 | 11.19 |
| GCF_014174405.1 | 0.00 | 0.4 | 0.0 | 20.38 | 20.90 | 1.23 |
| GCF_000210415.1 | 0.37 | 1.1 | 0.0 | 22.39 | 0.47 | 11.19 |
| GCF_000214 | 0.32 | 1.1 | 0.0 | 26.38 | 16.57 | 10.29 |

| | | | | | | |
|---|---|---|---|---|---|---|
| 455.1 | | | | | | |
| GCF_000007085.1 | 0.60 | 0.6 | 0.0 | 29.83 | 0.15 | 9.5 |
| GCF_900537995.1 | 1.57 | 0.0 | 0.0 | 20.60 | 0.19 | 15.54 |
| GCF_020991205.1 | 0.78 | 0.5 | 0.0 | 26.21 | 1.30 | 0.01 |
| GCF_017894345.1 | 0.00 | 0.5 | 0.0 | 23.63 | 0.44 | 0.09 |
| GCF_902386585.1 | 0.00 | 0.2 | 0.0 | 21.56 | 0.36 | 9.88 |
| GCF_900196735.1 | 0.00 | 0.7 | 0.0 | 4.43 | 0.01 | 0.95 |
| GCF_021091115.1 | 0.00 | 0.7 | 0.0 | 4.96 | 0.01 | 0.42 |
| GCF_020042225.1 | 0.09 | 0.7 | 0.0 | 20.26 | 0.15 | 0.41 |
| GCF_020042125.1 | 0.09 | 0.7 | 0.0 | 20.64 | 0.14 | 0.62 |
| GCF_019443925.1 | 0.00 | 0.2 | 0.0 | 20.81 | 0.45 | 0.53 |
| GCF_018987235.1 | 0.09 | 0.5 | 0.0 | 21.14 | 0.06 | 0.26 |
| GCF_018885325.1 | 0.09 | 0.2 | 0.0 | 20.64 | 0.22 | 0.48 |
| GCF_016767795.1 | 0.09 | 0.5 | 0.0 | 21.50 | 0.10 | 0.36 |
| GCF_016647595.1 | 0.00 | 0.7 | 0.0 | 21.95 | 0.07 | 0.14 |
| GCF_015709185.1 | 0.00 | 0.0 | 0.0 | 22.56 | 0.60 | 9.92 |
| GCF_015377785.1 | 0.00 | 0.7 | 0.0 | 4.56 | 0.01 | 0.56 |
| GCF_014656585.1 | 0.32 | 0.0 | 0.0 | 22.03 | 0.52 | 9.92 |
| GCF_013487905.1 | 0.19 | 0.5 | 0.0 | 19.48 | 0.11 | 0.06 |
| GCF_013456 | 0.00 | 0.5 | 0.0 | 20.67 | 0.15 | 0.6 |

| | | | | | | |
|---|---|---|---|---|---|---|
| 995.1 | | | | | | |
| GCF_011044 195.1 | 0.00 | 0.7 | 0.0 | 4.27 | 0.01 | 0.82 |
| GCF_009933 525.1 | 0.19 | 0.7 | 0.0 | 20.10 | 0.22 | 9.67 |
| GCF_009769 205.1 | 0.00 | 0.2 | 0.0 | 20.20 | 0.09 | 9.17 |
| GCF_009734 125.1 | 0.00 | 0.7 | 0.0 | 4.67 | 0.01 | 9.15 |
| GCF_009184 665.1 | 1.21 | 0.5 | 0.0 | 20.71 | 0.50 | 9.82 |
| GCF_006740 305.1 | 0.00 | 0.7 | 0.0 | 3.86 | 0.00 | 9.15 |
| GCF_004114 755.1 | 0.00 | 0.0 | 0.0 | 22.11 | 0.59 | 9.92 |
| GCF_003999 355.1 | 0.97 | 0.5 | 0.0 | 20.44 | 0.20 | 7.2 |
| GCF_003971 565.1 | 0.09 | 0.2 | 0.0 | 21.18 | 0.17 | 9.87 |
| GCF_003955 865.1 | 0.00 | 0.0 | 0.0 | 22.22 | 0.60 | 9.92 |
| GCF_003795 065.1 | 0.65 | 0.7 | 0.0 | 21.38 | 0.16 | 9.81 |
| GCF_003610 975.1 | 0.00 | 0.5 | 0.0 | 19.87 | 0.24 | 10.04 |
| GCF_003597 655.1 | 0.00 | 0.5 | 0.0 | 4.80 | 0.01 | 9.15 |
| GCF_003351 805.1 | 0.00 | 0.7 | 0.0 | 4.72 | 0.00 | 9.15 |
| GCF_002849 955.1 | 0.00 | 0.2 | 0.0 | 21.39 | 0.32 | 10.08 |
| GCF_002849 935.1 | 0.00 | 0.2 | 0.0 | 20.48 | 0.35 | 10.24 |
| GCF_002285 775.1 | 0.00 | 0.7 | 0.0 | 5.87 | 0.01 | 9.95 |
| GCF_002278 095.1 | 0.00 | 0.7 | 0.0 | 4.41 | 0.01 | 9.2 |
| GCF_002142 | 0.00 | 0.7 | 0.0 | 3.80 | 0.01 | 9.15 |

| | | | | | | |
|---|---|---|---|---|---|---|
| 575.1 | | | | | | |
| GCF_001953 135.1 | 0.00 | 0.5 | 0.0 | 4.54 | 0.01 | 9.15 |
| GCF_001888 965.1 | 0.00 | 0.7 | 0.0 | 4.65 | 0.02 | 9.19 |
| GCF_001888 925.1 | 0.65 | 2.0 | 0.0 | 2.92 | 0.01 | 9.15 |
| GCF_001888 905.1 | 0.65 | 1.0 | 0.0 | 5.74 | 0.01 | 9.95 |
| GCF_001746 265.1 | 0.00 | 0.2 | 0.0 | 21.02 | 0.43 | 9.88 |
| GCF_001702 095.1 | 0.00 | 0.2 | 0.0 | 21.56 | 0.36 | 9.88 |
| GCF_001469 775.1 | 0.08 | 1.0 | 0.0 | 4.11 | 0.00 | 9.15 |
| GCF_001006 025.1 | 0.00 | 0.2 | 0.0 | 19.75 | 0.35 | 10.06 |
| GCF_000525 715.1 | 0.32 | 0.0 | 0.0 | 21.14 | 0.32 | 9.1 |
| GCF_000466 885.3 | 0.09 | 0.7 | 0.0 | 20.48 | 0.13 | 9.98 |
| GCF_000422 165.1 | 0.00 | 0.2 | 0.0 | 19.40 | 0.30 | 10.15 |
| GCF_000214 785.1 | 1.21 | 0.5 | 0.0 | 20.08 | 0.56 | 9.87 |
| GCF_000189 515.1 | 1.62 | 1.2 | 0.0 | 22.24 | 0.49 | 10.08 |
| GCF_000182 835.1 | 0.00 | 0.7 | 0.0 | 4.96 | 0.01 | 9.19 |
| GCF_002970 935.1 | 1.13 | 0.0 | 0.0 | 38.02 | 0.28 | 9.77 |
| GCF_002970 915.1 | 1.13 | 0.0 | 0.0 | 36.85 | 0.28 | 9.77 |
| GCF_008694 205.1 | 0.65 | 0.7 | 0.0 | 20.22 | 2.03 | 8.82 |
| GCF_001874 605.1 | 0.64 | 1.5 | 0.0 | 29.15 | 0.00 | 10.19 |
| GCF_014058 | 0.00 | 0.5 | 0.0 | 27.19 | 0.32 | 9.18 |

| | | | | | | |
|---|---|---|---|---|---|---|
| 685.1 | | | | | | |
| GCF_004010015.1 | 0.00 | 1.0 | 0.0 | 25.96 | 0.48 | 9.15 |
| GCF_000498675.1 | 0.00 | 0.5 | 0.0 | 25.65 | 0.19 | 7.78 |
| GCF_000193435.2 | 0.00 | 0.2 | 0.0 | 22.72 | 1.70 | 10.02 |
| GCF_020887095.1 | 0.00 | 0.5 | 0.0 | 21.71 | 0.10 | 0.8 |
| GCF_002762335.1 | 0.00 | 0.0 | 0.0 | 32.59 | 0.00 | 7.24 |
| GCF_021432145.1 | 0.00 | 1.0 | 0.0 | 22.16 | 0.02 | 0.35 |
| GCF_020883435.1 | 0.00 | 1.0 | 0.0 | 22.06 | 0.02 | 0.34 |
| GCF_020149995.1 | 0.00 | 0.2 | 0.0 | 21.10 | 0.23 | 0.02 |
| GCF_013342945.1 | 0.00 | 0.7 | 0.0 | 22.04 | 0.05 | 9.28 |
| GCF_003952845.1 | 0.00 | 0.5 | 0.0 | 22.16 | 0.02 | 9.22 |
| GCF_003047065.1 | 1.52 | 1.7 | 0.0 | 21.88 | 0.01 | 9.28 |
| GCF_002706375.1 | 0.00 | 0.2 | 0.0 | 20.85 | 0.24 | 9.51 |
| GCF_002286215.1 | 0.00 | 0.7 | 0.0 | 22.00 | 0.04 | 9.28 |
| GCF_002224305.1 | 0.00 | 0.7 | 0.0 | 22.35 | 0.01 | 9.28 |
| GCF_000961015.1 | 0.00 | 0.2 | 0.0 | 20.23 | 0.34 | 10.04 |
| GCF_000934625.1 | 0.00 | 0.7 | 0.0 | 21.84 | 0.03 | 9.28 |
| GCF_000389675.2 | 0.00 | 0.7 | 0.0 | 22.04 | 0.04 | 9.28 |
| GCF_000194115.1 | 0.00 | 0.2 | 0.0 | 21.23 | 0.06 | 9.16 |
| GCF_000191 | 0.00 | 0.2 | 0.0 | 21.18 | 0.13 | 9.19 |

| | | | | | | |
|---|---|---|---|---|---|---|
| 545.1 | | | | | | |
| GCF_000011985.1 | 0.00 | 0.7 | 0.0 | 22.03 | 0.04 | 9.28 |
| GCF_021278925.1 | 0.48 | 1.7 | 0.0 | 20.48 | 1.64 | 1.53 |
| GCF_007570935.1 | 0.00 | 0.5 | 0.0 | 20.87 | 39.73 | 9.33 |
| GCF_001888945.1 | 0.00 | 0.7 | 0.0 | 3.62 | 0.00 | 9.15 |
| GCF_022213385.1 | 0.00 | 0.5 | 0.0 | 26.79 | 0.35 | 0.03 |
| GCF_021278945.1 | 0.09 | 0.2 | 0.0 | 21.10 | 0.20 | 0.31 |
| GCF_002849915.1 | 0.00 | 0.2 | 0.0 | 20.83 | 0.34 | 10.03 |
| GCF_004011315.1 | 0.39 | 0.7 | 0.0 | 27.52 | 0.13 | 9.2 |
| GCF_009730255.1 | 0.00 | 0.0 | 0.0 | 22.55 | 0.05 | 5.1 |
| GCF_001936235.1 | 0.00 | 0.2 | 0.0 | 23.36 | 0.19 | 7.22 |
| GCF_001308285.1 | 0.32 | 0.0 | 0.0 | 22.27 | 0.60 | 10.05 |
| GCF_900312975.1 | 0.00 | 0.0 | 0.0 | 26.83 | 0.09 | 1.05 |
| GCF_019597925.1 | 0.00 | 0.0 | 0.0 | 24.37 | 0.06 | 1.25 |
| GCF_014287355.1 | 0.00 | 0.4 | 0.0 | 24.63 | 35.82 | 10.61 |
| GCF_000145275.1 | 1.61 | 1.9 | 0.0 | 17.96 | 0.02 | 15.83 |
| GCF_021398395.1 | 0.00 | 0.2 | 0.0 | 21.29 | 5.50 | 0.28 |
| GCF_002075105.1 | 0.32 | 0.5 | 0.0 | 20.14 | 0.20 | 7.2 |
| GCF_000092405.1 | 0.51 | 0.8 | 0.0 | 26.17 | 0.01 | 10.16 |
| GCF_000686 | 0.40 | 0.0 | 0.0 | 20.06 | 21.52 | 10.8 |

| | | | | | | |
|---|---|---|---|---|---|---|
| 065.1 | | | | | | |
| GCF_000746025.2 | 1.91 | 1.4 | 0.0 | 27.58 | 0.00 | 10.66 |
| GCF_009730275.1 | 0.09 | 0.7 | 0.0 | 21.36 | 0.08 | 9.87 |
| GCF_009498395.1 | 0.00 | 0.2 | 0.0 | 19.30 | 0.31 | 10.06 |
| GCF_001888985.1 | 0.00 | 0.7 | 0.0 | 1.57 | 0.01 | 9.15 |
| GCF_000015385.1 | 0.00 | 0.5 | 0.0 | 20.86 | 0.46 | 9.88 |
| GCF_900155615.1 | 0.00 | 0.4 | 0.0 | 22.62 | 14.51 | 1.29 |
| GCF_022174665.1 | 0.00 | 0.0 | 0.0 | 21.87 | 0.04 | 1.05 |
| GCF_014467055.1 | 0.00 | 0.4 | 0.0 | 24.28 | 0.00 | 1.61 |
| GCF_013340725.1 | 0.00 | 0.8 | 0.0 | 28.99 | 0.09 | 8.58 |
| GCF_013315815.1 | 0.00 | 1.6 | 0.0 | 29.92 | 0.10 | 9.05 |
| GCF_006714885.1 | 1.36 | 2.4 | 0.0 | 24.75 | 21.38 | 13.44 |
| GCF_004154955.1 | 0.00 | 0.0 | 0.0 | 22.94 | 0.26 | 10.55 |
| GCF_002564155.1 | 0.00 | 0.0 | 0.0 | 21.95 | 1.05 | 10.56 |
| GCF_020892115.1 | 0.00 | 0.0 | 0.0 | 24.79 | 0.04 | 0.41 |
| GCF_000723465.1 | 0.00 | 0.0 | 0.0 | 20.06 | 0.05 | 10.62 |
| GCF_900461425.1 | 0.14 | 1.9 | 0.0 | 24.31 | 1.21 | 10.13 |
| GCF_900447145.1 | 0.14 | 1.9 | 0.0 | 24.22 | 1.05 | 0.0 |
| GCF_900187165.1 | 0.87 | 0.4 | 0.0 | 27.27 | 0.24 | 0.01 |
| GCF_019056 | 0.74 | 1.5 | 0.0 | 22.18 | 0.85 | 13.67 |

| | | | | | | |
|---|---|---|---|---|---|---|
| 555.1 | | | | | | |
| GCF_004006395.2 | 1.76 | 1.1 | 0.0 | 22.71 | 0.34 | 13.67 |
| GCF_003013635.1 | 0.10 | 1.9 | 0.0 | 25.01 | 0.16 | 10.13 |
| GCF_001484725.1 | 0.50 | 1.5 | 0.0 | 22.92 | 38.58 | 12.63 |
| GCF_000253195.1 | 0.00 | 0.8 | 0.0 | 22.62 | 0.54 | 10.7 |
| GCF_000143685.1 | 0.50 | 1.5 | 0.0 | 23.17 | 41.85 | 13.67 |
| GCF_000484505.1 | 0.50 | 1.5 | 0.0 | 22.82 | 38.96 | 12.63 |
| GCF_012317385.1 | 0.00 | 1.1 | 0.0 | 24.06 | 0.24 | 10.13 |
| GCF_010669305.1 | 1.06 | 0.4 | 0.0 | 19.34 | 0.00 | 10.61 |
| GCF_000018325.1 | 0.95 | 0.4 | 0.0 | 25.01 | 0.01 | 10.69 |
| GCF_000270305.1 | 1.72 | 0.0 | 0.0 | 25.00 | 0.03 | 10.29 |
| GCF_900446985.1 | 0.87 | 0.8 | 0.0 | 26.88 | 1.72 | 0.23 |
| GCF_008281175.1 | 0.00 | 0.4 | 0.0 | 23.67 | 0.00 | 13.24 |
| GCF_000305935.1 | 0.00 | 0.8 | 0.0 | 24.52 | 0.03 | 9.85 |
| GCF_000196455.1 | 1.40 | 1.1 | 0.0 | 27.29 | 0.01 | 10.56 |
| GCF_900452355.1 | 0.43 | 1.2 | 0.0 | 27.92 | 0.29 | 1.56 |
| GCF_013487865.1 | 0.00 | 0.5 | 0.0 | 26.94 | 0.23 | 0.04 |
| GCF_001714745.1 | 0.00 | 0.5 | 0.0 | 26.51 | 0.35 | 9.23 |
| GCF_000014425.1 | 0.00 | 0.5 | 0.0 | 27.04 | 0.36 | 9.15 |
| GCF_900070 | 1.68 | 0.0 | 0.0 | 25.38 | 0.08 | 0.36 |

| | | | | | | |
|---|---|---|---|---|---|---|
| 325.1 | | | | | | |
| GCF_900447065.1 | 0.00 | 0.4 | 0.0 | 22.86 | 0.38 | 0.29 |
| GCF_013407315.1 | 0.81 | 0.4 | 0.0 | 19.28 | 0.06 | 0.07 |
| GCF_013297165.1 | 1.61 | 1.9 | 0.0 | 19.00 | 0.09 | 1.05 |
| GCF_003628755.1 | 0.81 | 0.4 | 0.0 | 23.83 | 0.03 | 10.14 |
| GCF_002312985.1 | 0.00 | 0.8 | 0.0 | 22.37 | 0.54 | 10.13 |
| GCF_012848655.1 | 1.52 | 1.0 | 0.0 | 20.28 | 0.04 | 7.2 |
| GCF_008831485.1 | 0.87 | 0.2 | 0.0 | 21.04 | 0.03 | 7.2 |
| GCF_001042405.1 | 1.39 | 1.0 | 0.0 | 20.22 | 0.15 | 7.35 |
| GCF_000056065.1 | 0.00 | 0.5 | 0.0 | 4.44 | 0.01 | 9.15 |
| GCF_014467015.1 | 0.00 | 1.4 | 0.0 | 23.53 | 0.02 | 1.36 |
| GCF_000020005.1 | 1.69 | 1.2 | 0.0 | 23.05 | 0.00 | 10.52 |
| GCF_003057965.1 | 1.72 | 0.2 | 0.0 | 25.45 | 0.10 | 8.32 |
| GCF_000212395.1 | 1.72 | 0.4 | 0.0 | 25.40 | 0.03 | 8.82 |
| GCF_000007625.1 | 0.14 | 1.9 | 0.0 | 24.51 | 0.49 | 10.13 |
| GCF_016766995.1 | 0.50 | 1.5 | 0.0 | 22.27 | 0.29 | 0.41 |
| GCF_020892095.1 | 0.88 | 0.0 | 0.0 | 25.90 | 0.08 | 0.66 |
| GCF_000328625.1 | 0.00 | 1.2 | 0.0 | 27.01 | 0.03 | 9.84 |
| GCF_001940235.1 | 0.71 | 1.1 | 0.0 | 20.83 | 15.95 | 10.13 |
| GCF_003932 | 0.32 | 0.4 | 0.0 | 26.59 | 0.00 | 9.52 |

| | | | | | | |
|---|---|---|---|---|---|---|
| 015.1 | | | | | | |
| GCF_000018425.1 | 0.32 | 0.4 | 0.0 | 25.55 | 0.00 | 10.09 |
| GCF_000092965.1 | 0.24 | 0.0 | 0.0 | 30.81 | 1.75 | 9.83 |
| GCF_017301615.1 | 1.80 | 1.6 | 0.0 | 28.54 | 0.01 | 0.81 |
| GCF_003457035.1 | 0.67 | 1.1 | 0.0 | 21.96 | 0.32 | 11.19 |
| GCF_003457015.1 | 0.67 | 1.1 | 0.0 | 21.95 | 0.29 | 11.19 |
| GCF_003456975.1 | 0.67 | 1.1 | 0.0 | 21.89 | 0.34 | 11.19 |
| GCF_003697225.1 | 0.34 | 1.5 | 0.0 | 22.66 | 0.33 | 10.98 |
| GCF_003697205.1 | 0.34 | 1.5 | 0.0 | 22.59 | 0.36 | 10.98 |
| GCF_000009905.1 | 0.03 | 0.8 | 0.0 | 21.80 | 0.00 | 11.85 |
| GCF_000732725.1 | 1.63 | 0.8 | 0.0 | 20.85 | 16.97 | 15.54 |
| GCF_004103755.1 | 0.00 | 1.6 | 0.0 | 26.15 | 0.02 | 10.66 |
| GCF_902376065.1 | 0.81 | 0.8 | 0.0 | 26.43 | 18.91 | 0.46 |
| GCF_900087015.1 | 0.81 | 0.8 | 0.0 | 26.43 | 18.91 | 0.46 |
| GCF_002285495.1 | 1.24 | 1.5 | 0.0 | 25.63 | 0.20 | 9.76 |
| GCF_000213255.1 | 1.48 | 0.8 | 0.0 | 24.80 | 0.02 | 10.56 |
| GCF_900683755.1 | 0.16 | 0.8 | 0.0 | 19.40 | 0.05 | 15.74 |
| GCF_000243135.2 | 1.36 | 0.0 | 0.0 | 24.14 | 0.01 | 10.6 |
| GCF_003614235.1 | 0.69 | 1.1 | 0.0 | 23.85 | 36.30 | 9.77 |
| GCF_000163 | 0.70 | 1.9 | 0.0 | 27.12 | 0.40 | 7.39 |

| | | | | | | |
|---|---|---|---|---|---|---|
| 895.2 | | | | | | |
| GCF_900683775.1 | 0.00 | 0.8 | 0.0 | 20.78 | 0.06 | 15.66 |
| GCF_000688015.1 | 0.35 | 0.4 | 0.0 | 18.82 | 14.46 | 8.87 |
| GCF_014303955.1 | 1.68 | 1.6 | 0.0 | 23.16 | 0.00 | 2.16 |
| GCF_009738535.1 | 0.65 | 0.2 | 0.0 | 22.61 | 0.10 | 7.35 |
| GCF_003173695.1 | 1.14 | 0.5 | 0.0 | 18.55 | 0.64 | 7.78 |
| GCF_000761135.1 | 1.14 | 0.5 | 0.0 | 18.57 | 0.39 | 7.78 |
| GCF_900095895.1 | 0.93 | 2.3 | 0.0 | 23.32 | 0.47 | 3.09 |
| GCF_018603475.1 | 0.37 | 1.5 | 0.0 | 23.35 | 0.27 | 0.75 |
| GCF_018603455.1 | 0.37 | 1.5 | 0.0 | 23.26 | 0.26 | 0.75 |
| GCF_018603395.1 | 0.37 | 1.5 | 0.0 | 23.28 | 0.28 | 0.75 |
| GCF_016767135.1 | 1.99 | 1.9 | 0.0 | 22.36 | 0.28 | 0.66 |
| GCF_016767115.1 | 1.83 | 1.5 | 0.0 | 22.56 | 0.28 | 10.98 |
| GCF_016767095.1 | 1.42 | 3.4 | 0.0 | 22.85 | 0.40 | 2.95 |
| GCF_016767055.1 | 0.99 | 1.5 | 0.0 | 22.86 | 0.39 | 10.98 |
| GCF_016767035.1 | 0.99 | 1.5 | 0.0 | 22.35 | 0.35 | 0.47 |
| GCF_016767015.1 | 0.99 | 1.5 | 0.0 | 22.22 | 0.30 | 1.43 |
| GCF_016766975.1 | 1.47 | 1.9 | 0.0 | 22.91 | 0.37 | 1.36 |
| GCF_016766955.1 | 0.69 | 1.5 | 0.0 | 23.47 | 0.43 | 1.17 |
| GCF_003482 | 0.53 | 1.5 | 0.0 | 22.95 | 0.39 | 10.98 |

| | | | | | | |
|---|---|---|---|---|---|---|
| 365.1 | | | | | | |
| GCF_003482165.1 | 0.53 | 1.5 | 0.0 | 23.18 | 0.34 | 11.19 |
| GCF_002946555.2 | 0.53 | 1.5 | 0.0 | 23.06 | 0.34 | 10.98 |
| GCF_002946535.2 | 0.53 | 1.5 | 0.0 | 22.87 | 0.35 | 10.98 |
| GCF_002946515.2 | 0.53 | 1.5 | 0.0 | 23.01 | 0.34 | 10.98 |
| GCF_001972115.1 | 0.34 | 1.5 | 0.02 | 22.97 | 0.39 | 10.98 |
| GCF_018884725.1 | 0.50 | 1.5 | 0.0 | 22.46 | 0.25 | 0.51 |
| GCF_003287895.1 | 0.32 | 0.4 | 0.0 | 25.04 | 0.14 | 10.66 |
| GCF_000210435.1 | 0.37 | 1.5 | 0.0 | 23.07 | 0.42 | 10.98 |
| GCF_000144695.1 | 0.00 | 0.8 | 0.0 | 28.70 | 0.02 | 9.91 |
| GCF_019469365.1 | 0.84 | 0.5 | 0.0 | 19.91 | 0.00 | 9.19 |
| GCF_019469345.1 | 1.58 | 0.5 | 0.0 | 18.79 | 0.01 | 1.5 |
| GCF_019469325.1 | 1.49 | 0.2 | 0.0 | 19.89 | 0.00 | 0.87 |
| GCF_019469265.1 | 1.49 | 0.5 | 0.0 | 20.01 | 0.01 | 1.11 |
| GCF_003202825.1 | 1.62 | 0.2 | 0.0 | 18.72 | 27.96 | 9.29 |
| GCF_003151025.1 | 0.97 | 0.5 | 0.0 | 18.87 | 0.04 | 9.18 |
| GCF_000014725.1 | 0.21 | 1.5 | 0.0 | 23.19 | 0.00 | 10.38 |
| GCF_003150935.1 | 0.76 | 0.5 | 0.0 | 19.31 | 0.00 | 5.54 |
| GCF_015140235.1 | 0.00 | 0.0 | 0.0 | 24.62 | 0.04 | 1.24 |
| GCF_005222 | 0.34 | 0.0 | 0.0 | 23.79 | 0.01 | 10.45 |

| | | | | | | |
|---|---|---|---|---|---|---|
| 505.1 | | | | | | |
| GCF_005121165.2 | 0.00 | 0.0 | 0.0 | 21.21 | 42.86 | 14.17 |
| GCF_003612855.1 | 0.34 | 0.0 | 0.0 | 23.55 | 0.01 | 10.52 |
| GCF_003612835.1 | 0.34 | 0.0 | 0.0 | 24.08 | 0.01 | 10.45 |
| GCF_003020045.1 | 0.34 | 0.0 | 0.0 | 23.77 | 0.01 | 10.45 |
| GCF_000178115.2 | 0.34 | 0.0 | 0.0 | 23.65 | 0.01 | 10.45 |
| GCF_009734445.1 | 1.06 | 0.4 | 0.0 | 25.56 | 0.02 | 10.11 |
| GCF_000213235.1 | 0.96 | 1.3 | 0.0 | 28.99 | 0.00 | 10.06 |
| GCF_000328765.2 | 0.96 | 1.1 | 0.0 | 28.87 | 0.00 | 10.06 |
| GCF_001561955.1 | 0.00 | 1.1 | 0.0 | 21.87 | 0.00 | 10.58 |
| GCF_010120715.1 | 0.35 | 1.5 | 0.0 | 17.98 | 0.01 | 10.66 |
| GCF_015159595.1 | 0.00 | 0.4 | 0.0 | 25.31 | 0.00 | 10.85 |
| GCF_006408235.1 | 0.37 | 1.5 | 0.0 | 22.64 | 0.36 | 11.19 |
| GCF_010669085.1 | 0.35 | 0.4 | 0.0 | 26.13 | 0.06 | 9.5 |
| GCF_003600355.1 | 1.38 | 0.4 | 0.0 | 26.26 | 0.06 | 10.13 |
| GCF_013363915.1 | 0.00 | 0.5 | 0.0 | 23.80 | 18.35 | 0.34 |
| GCF_000144645.1 | 0.00 | 0.2 | 0.0 | 30.42 | 0.00 | 9.75 |
| GCF_000955745.1 | 1.11 | 0.6 | 0.0 | 23.74 | 43.82 | 10.37 |
| GCF_018919205.1 | 0.67 | 0.0 | 0.0 | 22.31 | 7.18 | 12.1 |
| GCF_003718 | 1.61 | 0.8 | 0.0 | 25.98 | 0.07 | 10.13 |

| | | | | | | |
|---|---|---|---|---|---|---|
| 715.1 | | | | | | |
| GCF_000025 225.2 | 0.71 | 0.0 | 0.0 | 27.62 | 0.14 | 5.19 |
| GCF_003017 335.1 | 0.00 | 1.1 | 0.0 | 23.01 | 24.47 | 10.98 |
| GCF_018408 575.1 | 0.67 | 0.4 | 0.0 | 23.26 | 0.10 | 0.47 |

**Table S1: Level of contamination estimation in reference genomes for the six tools**

| type | rank | # simulations |
|---|---|---|
| rate0-all | phylum | 92 |
| rate0-all | class | 133 |
| rate0-all | order | 114 |
| rate0-all | family | 132 |
| rate0-all | genus | 117 |
| rate0-all | species | 89 |
| rate10-all | phylum | 38 |
| rate10-all | class | 109 |
| rate10-all | order | 125 |
| rate10-all | family | 79 |
| rate10-all | genus | 107 |
| rate10-all | species | 44 |
| rate25-all | phylum | 63 |
| rate25-all | class | 52 |
| rate25-all | order | 137 |
| rate25-all | family | 87 |
| rate25-all | genus | 134 |
| rate25-all | species | 100 |
| rate0-redundant | phylum | 67 |

| | | |
|---|---|---|
| rate0-redundant | class | 114 |
| rate0-redundant | order | 108 |
| rate0-redundant | family | 115 |
| rate0-redundant | genus | 120 |
| rate0-redundant | species | 153 |
| rate0-replacement | phylum | 83 |
| rate0-replacement | class | 110 |
| rate0-replacement | order | 129 |
| rate0-replacement | family | 52 |
| rate0-replacement | genus | 126 |
| rate0-replacement | species | 153 |
| rate0-single | phylum | 106 |
| rate0-single | class | 121 |
| rate0-single | order | 118 |
| rate0-single | family | 66 |
| rate0-single | genus | 138 |
| rate0-single | species | 107 |

**Table S2: Number of simulations used.**

| trial | rank | redundant level | replacement level | single level | median simulation |
|---|---|---|---|---|---|
| rate0-all | phylum | 2.0 | 2.0 | 4.0 | 4.811932623820972 |
| rate0-all | class | 2.0 | 2.0 | 4.0 | 4.873136198540901 |
| rate0-all | order | 2.0 | 2.0 | 4.0 | 5.019076210557307 |
| rate0-all | family | 3.0 | 3.0 | 4.0 | 6.1572178296457345 |
| rate0-all | genus | 4.0 | 4.0 | 3.0 | 6.577107768849444 |
| rate0-all | species | 4.0 | 4.0 | 2.0 | 6.13509647963616 |
| rate10-all | phylum | 2.0 | 2.0 | 4.0 | 4.834503241391079 |
| rate10-all | class | 2.0 | 2.0 | 4.0 | 4.878188356692574 |
| rate10-all | order | 2.0 | 2.0 | 4.0 | 4.846119892534356 |

| | | | | | |
|---|---|---|---|---|---|
| rate10-all | family | 3.0 | 3.0 | 4.0 | 6.3780520629573765 |
| rate10-all | genus | 4.0 | 4.0 | 3.0 | 6.756596677229247 |
| rate10-all | species | 4.0 | 2.0 | 4.0 | 5.846661435621631 |
| rate25-all | phylum | 2.0 | 2.0 | 4.0 | 4.908778572089017 |
| rate25-all | class | 3.0 | 3.0 | 4.0 | 6.272828793654588 |
| rate25-all | order | 2.0 | 2.0 | 4.0 | 5.017066148552706 |
| rate25-all | family | 3.0 | 3.0 | 4.0 | 5.839175682683982 |
| rate25-all | genus | 4.0 | 4.0 | 3.0 | 6.72012660353524 |
| rate25-all | species | 4.0 | 4.0 | 2.0 | 5.8049846004168515 |
| rate0-redundant | phylum | 4.0 | 0.0 | 0.0 | 2.6791541661854485 |
| rate0-redundant | class | 4.0 | 0.0 | 0.0 | 2.530434968221652 |
| rate0-redundant | order | 4.0 | 0.0 | 0.0 | 2.669250167328002 |
| rate0-redundant | family | 6.0 | 0.0 | 0.0 | 3.891735096323639 |
| rate0-duplication | genus | 8.0 | 0.0 | 0.0 | 5.2260866932698065 |
| rate0-redundant | species | 8.0 | 0.0 | 0.0 | 5.008258180235766 |
| rate0-replacement | phylum | 0.0 | 4.0 | 0.0 | 2.7405731208809545 |
| rate0-replacement | class | 0.0 | 4.0 | 0.0 | 2.851081057089025 |
| rate0-replacement | order | 0.0 | 4.0 | 0.0 | 2.8589954433991465 |
| rate0-replacement | family | 0.0 | 6.0 | 0.0 | 4.151663232543372 |
| rate0-replacement | genus | 0.0 | 8.0 | 0.0 | 5.361825783843343 |
| rate0-replacement | species | 0.0 | 8.0 | 0.0 | 5.2731246552877575 |
| rate0-single | phylum | 0.0 | 0.0 | 8.0 | 4.113109358121008 |
| rate0-single | class | 0.0 | 0.0 | 8.0 | 4.756249845972857 |
| rate0-single | order | 0.0 | 0.0 | 8.0 | 4.444640457488092 |

| | | | | | |
|---|---|---|---|---|---|
| rate0-single | family | 0.0 | 0.0 | 8.0 | 4.73017412136555 |
| rate0-single | genus | 0.0 | 0.0 | 3.0 | 1.6826151227745412 |
| rate0-single | species | 0.0 | 0.0 | 2.0 | 1.017128662251577 |

**Table S3: Chimeric levels of the simulations.**

| type | rank | tool | percentage of underdetection | median under detection | median simulation |
|---|---|---|---|---|---|
| rate0-all | phylum | CheckM | 2.1739130434782608 | 0.547367806281502 | 4.811932623820972 |
| rate0-all | phylum | BUSCO | 20.652173913043477 | 2.9391841151873885 | 4.811932623820972 |
| rate0-all | phylum | GUNC | 59.78260869565217 | 4.85858972795982 | 4.811932623820972 |
| rate0-all | phylum | Physeter | 30.434782608695656 | 2.43565650512042 | 4.811932623820972 |
| rate0-all | phylum | Kraken2 | 16.304347826086957 | 0.12644517470050687 | 4.811932623820972 |
| rate0-all | phylum | CheckM2 | 2.1739130434782608 | 0.34084134735905547 | 4.811932623820972 |
| rate0-all | class | CheckM | 3.007518796992481 | 1.341675015238512 | 4.873136198540901 |
| rate0-all | class | BUSCO | 8.270676691729323 | 0.8906496557598418 | 4.873136198540901 |
| rate0-all | class | GUNC | 65.41353383458647 | 4.852431908251488 | 4.873136198540901 |
| rate0-all | class | Physeter | 0 | NA | 4.873136198540901 |
| rate0-all | class | Kraken2 | 30.07518796992481 2 | 2.1689779719793023 | 4.873136198540901 |
| rate0-all | class | CheckM2 | 6.015037593984962 | 0.9509236442434488 | 4.873136198540901 |
| rate0-all | order | CheckM | 7.894736842105263 | 3.672384429609872 | 5.019076210557307 |
| rate0-all | order | BUSCO | 15.789473684210526 | 1.2571785767979746 | 5.019076210557307 |
| rate0-all | order | GUNC | 78.94736842105263 | 4.98685311620279 | 5.019076210557307 |

| | | | | | |
|---|---|---|---|---|---|
| rate0-all | order | Physeter | 0 | NA | 5.019076210557307 |
| rate0-all | order | Kraken2 | 13.157894736842104 | 0.0870568791469761З | 5.019076210557307 |
| rate0-all | order | CheckM2 | 13.157894736842104 | 0.6699853979044166 | 5.019076210557307 |
| rate0-all | family | CheckM | 0 | NA | 6.1572178296457345 |
| rate0-all | family | BUSCO | 3.787878787878788 | 0.8099278755693158 | 6.1572178296457345 |
| rate0-all | family | GUNC | 63.63636363636363 | 6.155475937044468 | 6.1572178296457345 |
| rate0-all | family | Physeter | 0 | NA | 6.1572178296457345 |
| rate0-all | family | Kraken2 | 15.90909090909090 8 | 0.48133315562017387 | 6.1572178296457345 |
| rate0-all | family | CheckM2 | 6.8181818181818175 | 1.216142742601419 | 6.1572178296457345 |
| rate0-all | genus | CheckM | 5.982905982905983 | 3.2880552800398446 | 6.577107768849444 |
| rate0-all | genus | BUSCO | 11.11111111111111 | 1.6897786638765142 | 6.577107768849444 |
| rate0-all | genus | GUNC | 100.0 | 6.56604700623439 9 | 6.577107768849444 |
| rate0-all | genus | Physeter | 2.564102564102564 | 1.6127173173577631 | 6.577107768849444 |
| rate0-all | genus | Kraken2 | 51.28205128205128 | 3.702722366008586 | 6.577107768849444 |
| rate0-all | genus | CheckM2 | 12.82051282051282 | 0.8280552800398446 | 6.577107768849444 |
| rate0-all | species | CheckM | 19.101123595505616 | 1.0230093681452876 | 6.13509647963616 |
| rate0-all | species | BUSCO | 33.70786516853933 | 2.966149927016671 | 6.13509647963616 |
| rate0-all | species | GUNC | 100.0 | 6.099406536092411 | 6.13509647963616 |
| rate0-all | species | Physeter | 2.247191011235955 | 1.1266853151578897 | 6.13509647963616 |

| | | | | | |
|---|---|---|---|---|---|
| rate0-all | species | Kraken2 | 97.75280898876404 | 5.708448938543611 | 6.13509647963616 |
| rate0-all | species | CheckM2 | 23.59550561797526 | 1.17896236484225 | 6.13509647963616 |
| rate0-redundant | phylum | CheckM | 0 | NA | 2.6791541661854485 |
| rate0-redundant | phylum | BUSCO | 10.44776119402985 | 0.8777160613884001 | 2.6791541661854485 |
| rate0-redundant | phylum | GUNC | 49.25373134328358 | 2.7680049896122947 | 2.6791541661854485 |
| rate0-redundant | phylum | Physeter | 32.83582089552239 | 0.2752726041623468 | 2.6791541661854485 |
| rate0-redundant | phylum | Kraken2 | 10.44776119402985 | 0.9663416075099855 | 2.6791541661854485 |
| rate0-redundant | phylum | CheckM2 | 0 | NA | 2.6791541661854485 |
| rate0-redundant | class | CheckM | 0.8771929824561403 | 0.0738489295303495 | 2.530434968221652 |
| rate0-redundant | class | BUSCO | 0.8771929824561403 | 2.0638489295303493 | 2.530434968221652 |
| rate0-redundant | class | GUNC | 63.1578947368421 | 2.518422922309708 | 2.530434968221652 |
| rate0-redundant | class | Physeter | 0 | NA | 2.530434968221652 |
| rate0-redundant | class | Kraken2 | 4.385964912280701 | 0.007164387608035838 | 2.530434968221652 |
| rate0-redundant | class | CheckM2 | 0 | NA | 2.530434968221652 |
| rate0-redundant | order | CheckM | 0.9259259259259258 | 0.6612448837864955 | 2.669250167328002 |
| rate0-redundant | order | BUSCO | 0.9259259259259258 | 0.08914031096218045 | 2.669250167328002 |
| rate0-redundant | order | GUNC | 62.03703703703704 | 2.6702495466883254 | 2.669250167328002 |
| rate0-redundant | order | Physeter | 0 | NA | 2.669250167328002 |
| rate0-redundant | order | Kraken2 | 13.88888888888889 | 0.12302928900511256 | 2.669250167328002 |

| | | | | | |
|---|---|---|---|---|---|
| rate0-redundant | order | CheckM2 | 0 | NA | 2.669250167328002 |
| rate0-redundant | family | CheckM | 0 | NA | 3.891735096323639 |
| rate0-redundant | family | BUSCO | 0 | NA | 3.891735096323639 |
| rate0-redundant | family | GUNC | 71.30434782608695 | 3.797801660687545 | 3.891735096323639 |
| rate0-redundant | family | Physeter | 0 | NA | 3.891735096323639 |
| rate0-redundant | family | Kraken2 | 4.3478260869565215 | 0.7405467250892177 | 3.891735096323639 |
| rate0-redundant | family | CheckM2 | 0 | NA | 3.891735096323639 |
| rate0-redundant | genus | CheckM | 0 | NA | 5.2260866932698065 |
| rate0-redundant | genus | BUSCO | 0.8333333333333334 | 1.527495927267415 | 5.2260866932698065 |
| rate0-redundant | genus | GUNC | 98.33333333333333 | 5.221863701029459 | 5.2260866932698065 |
| rate0-redundant | genus | Physeter | 3.3333333333333335 | 0.9145162254893897 | 5.2260866932698065 |
| rate0-redundant | genus | Kraken2 | 48.33333333333336 | 2.925491955486992 | 5.2260866932698065 |
| rate0-redundant | genus | CheckM2 | 0 | NA | 5.2260866932698065 |
| rate0-redundant | species | CheckM | 2.6143790849673203 | 1.6146680333416403 | 5.008258180235766 |
| rate0-redundant | species | BUSCO | 11.11111111111111 | 0.23635017980810868 | 5.008258180235766 |
| rate0-redundant | species | GUNC | 99.34640522875817 | 4.980161206953943 | 5.008258180235766 |
| rate0-redundant | species | Physeter | 1.3071895424836601 | 0.8846836918623215 | 5.008258180235766 |
| rate0-redundant | species | Kraken2 | 96.73202614379085 | 4.596433622837286 | 5.008258180235766 |
| rate0-redundant | species | CheckM2 | 1.3071895424836601 | 0.4133483116012959 | 5.008258180235766 |

| | | | | | |
|---|---|---|---|---|---|
| rate0-replacement | phylum | CheckM | 60.24096385542169 | 2.4228506140333854 | 2.7405731208809545 |
| rate0-replacement | phylum | BUSCO | 100.0 | 2.270097042264002 | 2.7405731208809545 |
| rate0-replacement | phylum | GUNC | 83.13253012048193 | 2.7411761428531634 | 2.7405731208809545 |
| rate0-replacement | phylum | Physeter | 24.09638554216867 6 | 0.36839986350913034 | 2.7405731208809545 |
| rate0-replacement | phylum | Kraken2 | 10.843373493975903 | 0.018144475806775695 | 2.7405731208809545 |
| rate0-replacement | phylum | CheckM2 | 100.0 | 2.488929284161979 | 2.7405731208809545 |
| rate0-replacement | class | CheckM | 76.36363636363637 | 2.672664125650027 | 2.851081057089025 |
| rate0-replacement | class | BUSCO | 77.27272727272727 | 2.146217830384028 | 2.851081057089025 |
| rate0-replacement | class | GUNC | 91.81818181818183 | 2.8609219677617013 | 2.851081057089025 |
| rate0-replacement | class | Physeter | 0 | NA | 2.851081057089025 |
| rate0-replacement | class | Kraken2 | 10.0 | 0.0292669341801 0551 | 2.851081057089025 |
| rate0-replacement | class | CheckM2 | 96.36363636363636 | 2.3204636993396153 | 2.851081057089025 |
| rate0-replacement | order | CheckM | 70.54263565891473 | 2.718348929830576 | 2.8589954433991465 |
| rate0-replacement | order | BUSCO | 100.0 | 2.088465997125148 | 2.8589954433991465 |
| rate0-replacement | order | GUNC | 92.24806201550388 | 2.8589954433991465 | 2.8589954433991465 |
| rate0-replacement | order | Physeter | 0 | NA | 2.8589954433991465 |
| rate0-replacement | order | Kraken2 | 16.27906976744186 | 0.3167954719175756 | 2.8589954433991465 |
| rate0-replacement | order | CheckM2 | 97.67441860465115 | 2.2534782952096934 | 2.8589954433991465 |
| rate0-replacement | family | CheckM | 100.0 | 3.777301014277268 7 | 4.151663232543372 |

| | | | | | |
|---|---|---|---|---|---|
| rate0-replacement | family | BUSCO | 100.0 | 3.537099035261738 | 4.151663232543372 |
| rate0-replacement | family | GUNC | 94.23076923076923 | 4.1614971120859305 | 4.151663232543372 |
| rate0-replacement | family | Physeter | 0 | NA | 4.151663232543372 |
| rate0-replacement | family | Kraken2 | 26.923076923076923 | 0.053355342473088596 | 4.151663232543372 |
| rate0-replacement | family | CheckM2 | 98.07692307692307 | 3.5289765857154545 | 4.151663232543372 |
| rate0-replacement | genus | CheckM | 100.0 | 5.13551635367378 | 5.361825783843343 |
| rate0-replacement | genus | BUSCO | 100.0 | 4.578141603203857 | 5.361825783843343 |
| rate0-replacement | genus | GUNC | 100.0 | 5.361825783843343 | 5.361825783843343 |
| rate0-replacement | genus | Physeter | 4.761904761904762 | 0.34935489267373 | 5.361825783843343 |
| rate0-replacement | genus | Kraken2 | 53.96825396825397 | 2.5090024296601556 | 5.361825783843343 |
| rate0-replacement | genus | CheckM2 | 100.0 | 4.642534208737596 | 5.361825783843343 |
| rate0-replacement | species | CheckM | 100.0 | 4.948345885778904 | 5.2731246552877575 |
| rate0-replacement | species | BUSCO | 100.0 | 4.282675557042158 | 5.2731246552877575 |
| rate0-replacement | species | GUNC | 100.0 | 5.2731246552877575 | 5.2731246552877575 |
| rate0-replacement | species | Physeter | 2.6143790849673203 | 0.8762394590079698 | 5.2731246552877575 |
| rate0-replacement | species | Kraken2 | 97.38562091503267 | 4.924367217932163 | 5.2731246552877575 |
| rate0-replacement | species | CheckM2 | 99.34640522875817 | 4.64342400971791 | 5.2731246552877575 |
| rate0-single | phylum | CheckM | 98.11320754716981 | 2.753690553138729 | 4.113109358121008 |
| rate0-single | phylum | BUSCO | 100.0 | 3.3615234871499187 | 4.113109358121008 |

| | | | | | |
|---|---|---|---|---|---|
| rate0-single | phylum | GUNC | 62.264150943396224 | 4.063274297657863 | 4.113109358121008 |
| rate0-single | phylum | Physeter | 24.528301886792452 | 2.459039899001997 | 4.113109358121008 |
| rate0-single | phylum | Kraken2 | 5.660377358490567 | 0.033283677383916466 | 4.113109358121008 |
| rate0-single | phylum | CheckM2 | 96.2264150943396 3 | 3.01100625585747 | 4.113109358121008 |
| rate0-single | class | CheckM | 100.0 | 3.3957506248593834 | 4.756249845972857 |
| rate0-single | class | BUSCO | 100.0 | 3.8280322020257986 | 4.756249845972857 |
| rate0-single | class | GUNC | 66.94214876033058 | 4.762033999079752 | 4.756249845972857 |
| rate0-single | class | Physeter | 0 | NA | 4.756249845972857 |
| rate0-single | class | Kraken2 | 22.31404958677686 | 2.330137377753667 | 4.756249845972857 |
| rate0-single | class | CheckM2 | 99.17355371900827 | 3.67103717395642 | 4.756249845972857 |
| rate0-single | order | CheckM | 100.0 | 3.3533254432998714 | 4.444640457488092 |
| rate0-single | order | BUSCO | 100.0 | 3.577603145395972 | 4.444640457488092 |
| rate0-single | order | GUNC | 69.49152542372882 | 4.49297240404319 | 4.444640457488092 |
| rate0-single | order | Physeter | 0 | NA | 4.444640457488092 |
| rate0-single | order | Kraken2 | 19.491525423728813 | 0.0454989284027536 | 4.444640457488092 |
| rate0-single | order | CheckM2 | 96.61016949152543 | 3.585201776120518 6 | 4.444640457488092 |
| rate0-single | family | CheckM | 100.0 | 3.8815283203406183 | 4.73017412136555 |
| rate0-single | family | BUSCO | 100.0 | 4.027079880569484 | 4.73017412136555 |
| rate0-single | family | GUNC | 89.39393939393939 | 4.511252493568201 | 4.73017412136555 |

| | | | | | |
|---|---|---|---|---|---|
| rate0-single | family | Physeter | 0 | NA | 4.73017412136555 |
| rate0-single | family | Kraken2 | 16.666666666666664 | 0.5812524935682011 | 4.73017412136555 |
| rate0-single | family | CheckM2 | 100.0 | 3.892943683145122 | 4.73017412136555 |
| rate0-single | genus | CheckM | 91.30434782608695 | 1.341065010205717 | 1.6826151227745412 |
| rate0-single | genus | BUSCO | 88.40579710144928 | 0.8779338870099609 | 1.6826151227745412 |
| rate0-single | genus | GUNC | 94.20289855072464 | 1.6586791330048318 | 1.6826151227745412 |
| rate0-single | genus | Physeter | 0 | NA | 1.6826151227745412 |
| rate0-single | genus | Kraken2 | 33.33333333333333 | 0.7395275004820205 | 1.6826151227745412 |
| rate0-single | genus | CheckM2 | 92.02898550724638 | 1.1865059793997539 | 1.6826151227745412 |
| rate0-single | species | CheckM | 81.30841121495327 | 0.9039737079849057 | 1.017128662251577 |
| rate0-single | species | BUSCO | 57.943925233644855 | 0.6150415134750611 | 1.017128662251577 |
| rate0-single | species | GUNC | 97.19626168224299 | 1.0152493692973965 | 1.017128662251577 |
| rate0-single | species | Physeter | 0 | NA | 1.017128662251577 |
| rate0-single | species | Kraken2 | 85.04672897196261 | 0.6839737079849058 | 1.017128662251577 |
| rate0-single | species | CheckM2 | 73.83177570093457 | 0.6048218745330745 | 1.017128662251577 |

**Table S4: Underdetection of the tools.**
